# White matter microstructure predicts effort and reward sensitivity

**DOI:** 10.1101/2025.08.19.671080

**Authors:** Nam Trinh, Laurence Dricot, Pierre Vassiliadis, Quentin Dessain, Julie Duque, Tomas Ward, Gerard Derosiere

## Abstract

From rodents to humans, animals constantly face a central question: is the reward worth the effort? Effort and reward sensitivity in such situations vary substantially across individuals and ultimately shape goal-directed behavior. Yet, the brain mechanisms underlying this variability across individuals remain unclear. Here, we combined computational modeling of effort and reward sensitivity during decision-making with whole-brain diffusion MRI in 45 healthy participants to identify white matter substrates of individual sensitivity. A data-driven, cluster-based analysis of fractional anisotropy and mean diffusivity revealed 12 clusters: five linked to effort sensitivity, all within tracts connected to major frontal valuation nodes (e.g., supplementary motor area [SMA], dorsal anterior cingulate cortex [dACC], orbitofrontal cortex [OFC]), and seven linked to reward sensitivity, spanning frontal valuation, fronto-parietal, and sensorimotor networks. The strongest associations involved two SMA-connected clusters, one shared across effort and reward sensitivity and another consistent across both microstructural metrics. Critically, microstructural features from the five effort-related and seven reward-related clusters reliably predicted individual effort and reward sensitivity in out-of-sample machine learning analyses, respectively, whereas randomly sampled clusters did not. SMA-connected tracts were the dominant predictors in these decoding analyses, with additional contributions from fronto-parietal and sensorimotor pathways for reward sensitivity. These findings reveal a distributed white matter architecture underlying inter-individual differences in effort and reward sensitivity, with SMA pathways emerging as central hubs. They demonstrate that localized white matter microstructure can robustly predict these individual differences, offering a framework to forecast the impact of lesions or interventions on goal-directed behavior, including apathy and impulsivity.

**SIGNIFICANCE STATEMENT:** Why do some people give up easily when faced with high effort demands, while others persist even when rewards are small? Such differences in effort and reward sensitivity shape goal-directed behavior, yet their neural basis is unclear. Using diffusion MRI and computational modeling, we show that white matter microstructure in specific pathways reliably predicts individual differences in these sensitivities. Tracts connected to the supplementary motor area emerged as central hubs, with additional contributions from fronto-parietal and sensorimotor networks. These results demonstrate that variability in effort and reward sensitivity is rooted not only in brain activity but also in structural connectivity, providing a framework to anticipate how white matter lesions or interventions may alter goal-directed behavior, including apathy and impulsivity.

## INTRODUCTION

From rodents to humans, animals constantly face a central question: is the reward worth the effort? Everyday choices – doing chores or watching TV, cooking or ordering fast food – hinge on evaluating effort versus reward (Hogan et al., 2020). The drive to act in these situations depends on one’s sensitivity to each. Abnormal sensitivities disrupt goal-directed behavior: effort hypersensitivity leads to inaction despite potential rewards, contributing to apathy (Husain and Roiser, 2018; Costello et al., 2024), while reward hypersensitivity can drive action despite effort costs, fuelling impulsivity (Long et al., 2022; Luijten et al., 2017). Even in health, sensitivities vary widely (Bonnelle et al., 2016; Fuentes-Claramonte et al., 2016), shaping goal-directed behavior. These inter-individual differences – sometimes termed “computational phenotypes” (Pessiglione et al., 2018; Schurr et al., 2024) – can be captured by computational models of decision-making. Yet, the brain mechanisms behind this variability remain unclear. Identifying their anatomical basis could explain this variability and enable prediction of effort and reward sensitivity from brain structure (Thiebaut de Schotten and Forkel, 2022), offering a framework to forecast the impact of structural lesions on goal-directed behavior.

So far, insights into effort and reward processing have come from functional neuroimaging studies tracking brain activity across varying effort and reward levels during decision-making (Husain and Roiser, 2018). These highlight a core fronto- striatal network with partially dissociable roles: the supplementary motor area (SMA) tracks preferentially effort (Bonnelle et al., 2016), the orbitofrontal cortex (OFC) tracks reward (Klein-Flügge et al., 2022), while the dorsal anterior cingulate cortex (dACC; Shenhav et al., 2013) and nucleus accumbens (NAcc; Suzuki et al., 2021) encode both, integrating them into net value signals. Beyond this core, broader fronto-parietal (Etzel et al., 2016) and sensorimotor structures – including the cerebellum (Kostadinov and Häusser, 2022) and primary motor cortex (M1; Derosiere et al., 2025) – also show reward-related activity during decision-making. Together, these findings suggest that effort-reward choices rely on computations across distributed regions, with the fronto- striatal valuation network at the core.

What remains poorly understood is how individual differences in effort and reward sensitivity emerge from variability across this distributed system. While most studies focus on gray matter activity, growing evidence shows white matter also actively modulate, rather than just relay, neural signals, shaping behavior (Innocenti et al., 2022). Because effort and reward computations require coordination across many regions, structural connectivity may determine how these signals are processed (Thiebaut de Schotten and Forkel, 2022). This hypothesis has been rarely tested, and mainly using tract-of-interest approaches, as in our own work (Derosiere et al., 2024). For instance, subclinical apathy, linked to effort hypersensitivity, relates to anterior cingulum alterations, a tract connecting SMA, OFC and dACC (Bonnelle et al., 2016). More recently, we found that heightened effort sensitivity associates with reduced SMA-NAcc macroscopic connectivity (Derosiere et al., 2024). These findings suggest that white matter in tracts connecting fronto-striatal valuation regions shapes individual differences in effort and reward sensitivity. Still, given the wide distribution of value-related activity, whole-brain, data-driven approaches are now essential to determine the structural architecture underlying these individual differences.

Advances in diffusion imaging can help bridge this gap. Voxel-wise microstructure metrics, fractional anisotropy (FA) and mean diffusivity (MD), capture complementary features of axonal organization: FA reflects alignment, density, and myelination, whereas MD indexes extracellular space and reduced barriers (Beck et al., 2021; Song et al., 2025). Both are amenable to data-driven analyses, allowing identification of white matter loci with microstructural differences in one or both metrics. Here, we combined these analyses with computational modelling of effort and reward sensitivity during decision-making in 45 healthy participants to identify white matter regions whose microstructure covaries with, and predicts, sensitivity. A cluster-based approach unconstrained by anatomical priors revealed 12 clusters: 5 linked to effort sensitivity, all within tracts connected to frontal valuation nodes (SMA, dACC, OFC), and 7 linked to reward sensitivity, distributed across frontal valuation, fronto-parietal, and sensorimotor pathways. The strongest effects involved two SMA-connected clusters, one shared across effort and reward sensitivity and another consistent across FA and MD. Microstructural metrics from these clusters reliably predicted individual sensitivity in out-of-sample machine learning analyses, whereas randomly sampled clusters did not. Clusters in SMA-connected tracts were the dominant predictors in machine learning analyses, but fronto-parietal and sensorimotor pathways also contributed to the decoding of reward sensitivity. These findings reveal distributed white matter substrates for individual differences in effort and reward sensitivity and show that these sensitivities can be reliably decoded from localized microstructural features, offering a potential avenue to forecast the impact of structural lesions in patients with abnormal goal-directed behaviour, such as apathy and impulsivity.

## METHODS

### Participants

Fifty healthy adult participants were initially recruited from the Research Participant Pool of the Institute of Neuroscience at the Université catholique de Louvain (Brussels, Belgium). Three participants did not complete the full experimental protocol, and tractography could not be performed in two subjects due to corrupted MRI data. The final sample comprised 45 individuals (mean age = 25.1 ± 0.8 years; 31 females, 14 males), whose data were included in all subsequent analyses. All subjects were right-handed based on the Edinburgh Questionnaire (Oldfield, 1971) and had no history of neurological disorders, psychiatric illness, substance abuse, or use of medications that could affect performance. Participants received financial compensation for their involvement and could earn additional rewards based on task performance (see *Task description* section below). The study protocol was approved by the institutional review board of UCLouvain, and written informed consent was obtained from all participants.

A previous publication using this dataset of 45 participants investigated the relationship between apathy, decision-making and structural connectivity with a tract-of-interest approach, focusing on a restricted set of motor-related circuits, together with effective connectivity measures derived from transcranial magnetic stimulation (Derosiere, et al., 2024). That study relied on streamline-based tractography, which quantifies the reconstructed fiber tracts (streamlines) connecting two regions and reflects the macroscopic “size” or “strength” of a connection, but is largely insensitive to microstructural properties, and is focused on a limited set of tracts. In contrast, the present work addresses a distinct and complementary question by moving beyond predefined tracts to a whole-brain, data-driven framework. We leverage voxel-wise microstructural metrics (i.e., FA and MD) to capture complementary aspects of white matter integrity across the brain, and combine these with machine learning to identify the anatomical predictors of inter-individual variability in effort and reward sensitivity.

### Data acquisition

The data were acquired in two sessions, conducted with a minimum of interval of 24 hours and a maximum of interval of one week. In the first session, MRI data were acquired at the Saint-Luc University Hospitals (Brussels, Belgium). The second session, conducted at the Institute of Neuroscience, UCLouvain (Brussels), included behavioral assessment using an effort-based decision-making task commonly used in the field and allowing computational modelling of effort and reward sensitivities (Morris et al., 2025; Gilmour et al., 2023; Pessiglione et al., 2018). Detailed procedures for each session are described below.

### MRI data acquisition

Both structural T1-weighted and diffusion-weighted MRI data were acquired for each participant on a 3 Tesla MRI (SIGNA_TM_ Premier, General Electric), equipped with a 48-channel head coil (Figure 1.A). 3D T1-weighted anatomical images were obtained with the following parameters: Echo Time (TE) = 2.96 ms, Repetition Time (TR) = 2238.93 ms, Inversion Time TI = 900 ms, 170 slices, slice thickness = 1 mm, in-plane FOV = 256 × 256 mm², matrix = 256 × 256; voxel size = 1 mm_3_ isotropic. Diffusion-weighted MRI (DWIs) were acquired in the axial plane with the parameters: TR = 7289 ms, TE = 57.1 ms, 70 slices, slice thickness = 2 mm, in-plane FOV = 220 × 220 mm², matrix size = 110 × 110; 2 mm isotropic voxels, with 64 gradients at b = 1000 s/mm² and one b0 reference image.

**Figure 1:**
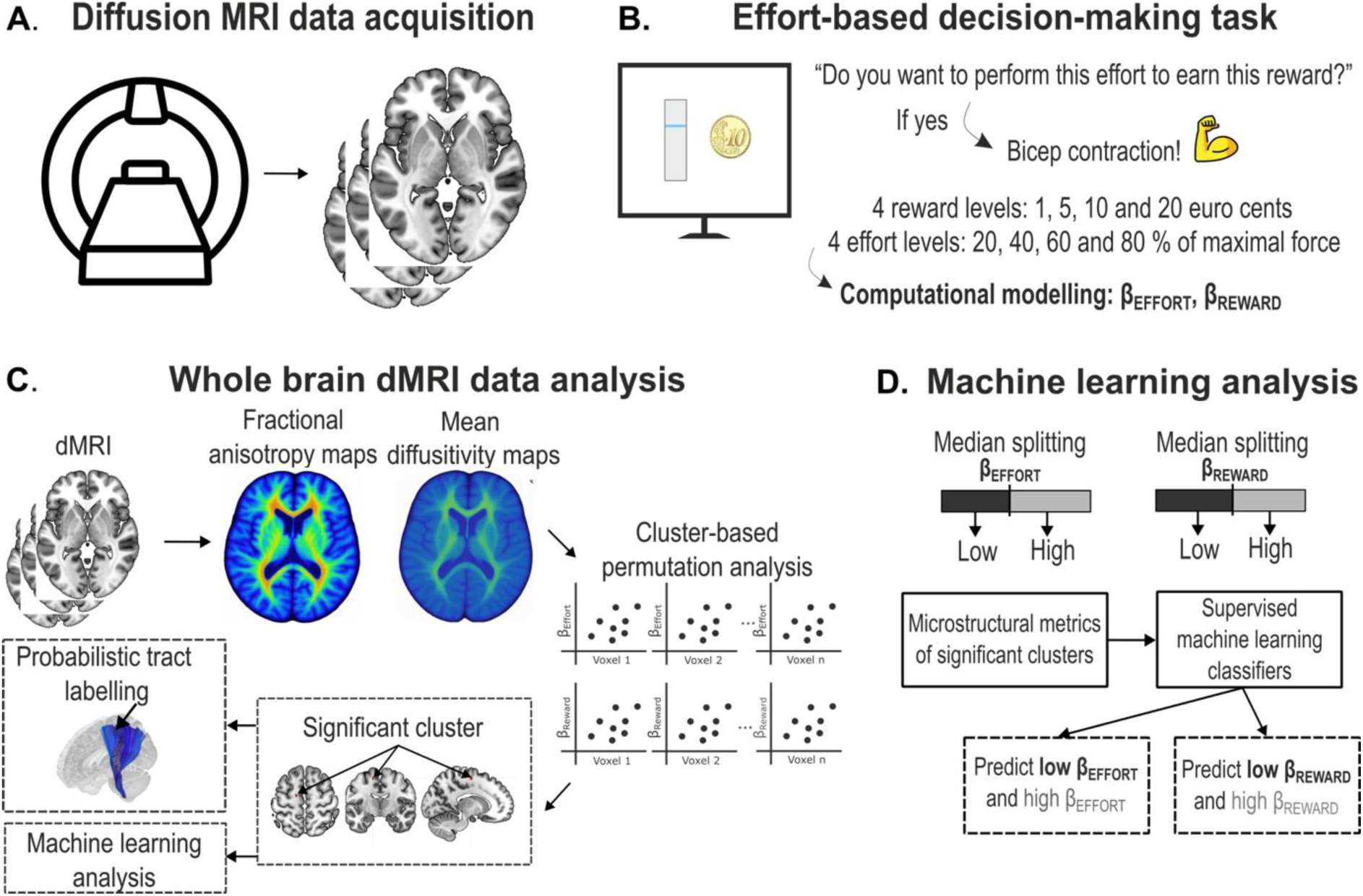
Experimental protocol and data analyses. **A. Diffusion MRI (dMRI) data acquisition.** dMRI data were acquired to assess white matter microstructure, including FA and MD. **B. Effort-based decision-making task and computational modelling of behavior.** Participants completed an effort-based decision-making task, choosing whether to perform biceps contractions of varying effort levels (20%, 40%, 60%, or 80% of maximal voluntary contraction) to obtain varying monetary rewards (1, 5, 10, or 20 euro cents). Computational modelling of their acceptance rates in this task, yielded individual effort and reward sensitivity parameters (β_Effort_ and β_Reward_, respectively). **C. Whole-brain dMRI data analysis.** dMRI data were exploited to compute voxel-wise FA and MD quantification in each subject (top left panel). We then applied a cluster-based analysis to identify significant clusters where FA and MD covaried with β_Effort_ or β_Reward_ (left and bottom panels). Significant clusters were mapped onto white matter tracts and characterized by MNI coordinates and their metrics were used in a predictive machine learning analysis (right panel; see D.). **D. Machine learning analysis.** Identified microstructural features were used as inputs for machine learning classifiers predicting low and high β_Effort_ and β_Reward_. Classifier performance was assessed using 5-fold cross-validation, yielding average accuracy and area under the receiving operator curve metrics.

### Behavioral data acquisition

#### Experiment setup

Participants engaged in an effort-based decision-making task on a computer while seated in an ergonomic chair positioned 100 cm from a monitor (refresh rate: 100 Hz). The right arm was flexed to 90 degrees and stabilized on an armrest, with participants gripping a custom-designed handle using the right hand. The left hand remained unrestrained and was used to make finger responses via the left and right arrow keys on a standard keyboard positioned on a table. To minimize movement artifacts and ensure consistent biomechanical positioning during force exertion, the right forearm was secured to the armrest using an adjustable strap.

#### Task description

The effort-based decision-making task was programmed using custom-written MATLAB scripts incorporating the Psychtoolbox library (Brainard, 1997). During this task, participants were required to evaluate whether to engage in varying levels of physical effort in exchange for varying monetary rewards (Figure 1.B). Effort was operationalized as an isometric contraction of the right biceps, elicited by instructing participants to attempt to flex their arm — bringing the fist toward the shoulder — while maintaining elbow contact with the armrest and gripping a fixed handle. The intensity of muscular contraction was continuously monitored via real-time surface electromyography. Monetary rewards were expressed in euro cents; participants were informed they would obtain the total amount earned during the task at the end of the session.

Each trial started with the display of a vertical force gauge accompanied by a horizontal reference line indicating the required force level for that trial. Adjacent to the gauge, the monetary reward was displayed. The full scale of the gauge represented 100% of the participant’s maximal voluntary contraction (MVC), which was individually calibrated prior to the experimental blocks (see *Blocks of trials and MVC* section, for details). Effort levels were set at 20%, 40%, 60%, or 80% of MVC, paired with potential rewards of 1, 5, 10, or 20 euro cents, resulting in 16 possible effort-reward combinations. Participants had 5 seconds to decide whether to accept or reject each offer, responding with their left index and middle fingers by pressing the right arrow key to accept or the left arrow key to reject.

Upon acceptance of an offer, a "Go" signal was presented on the screen following a variable delay of 1.0 to 1.2 seconds, marking the onset of the contraction period. During this period, participants received continuous visual biofeedback of their exertion level, depicted as a dynamic filling of the gauge. The extent of gauge filling was proportional to the rectified amplitude of the EMG signal recorded from the right biceps. Participants were instructed to maintain a contraction at or above the target force level — corresponding to 20%, 40%, 60%, or 80% of their MVC — for a minimum duration of 2.7 seconds within a 4-second window (i.e., >66% of the contraction period).

At the end of each trial, participants received visuo-auditory feedback indicating the outcome of the trial (success or failure) and whether the corresponding reward had been earned. Trials in which muscle activity was detected prior to the onset of the "Go" signal were immediately aborted, and an "Anticipated" message was displayed on the screen. Regardless of the outcome, a fixed 2-second inter-trial interval followed each trial to allow for recovery.

If participants declined the offer, the trial terminated immediately and was followed by a 2-second blank screen before the onset of the next trial (*i.e.*, corresponding to the inter-trial interval). Similarly, if no response was recorded within the initial 5-second decision window, the trial was considered a missed response, and the subsequent trial started after the standard inter-trial interval.

### Data analyses

#### Computational modelling of acceptance rates

To quantify individual differences in effort and reward sensitivity, we applied computational modelling to the acceptance rates in the effort-based decision-making task. As such, for each participant, the model estimated the influence of effort and reward on offer valuation, which was translated into a probability to accept the offer and engage in the action.

First, we tested a set of candidate models of subjective value computation informed by prior literature (Morris et al., 2025; Gilmour et al., 2023; Le Heron et al., 2018; Bonnelle et al., 2016), evaluating model fits using Bayesian information criterion (BIC) minimization and visual inspection (Supplementary Figure 1).

In the best-fitting model, the subjective value V of each offer was computed as:

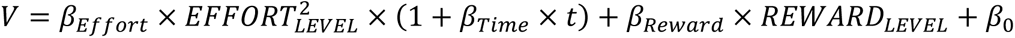

Here, the subjective value of an offer was modelled as a quadratic function of the proposed EFFORT_LEVEL_, and a linear function of the REWARD_LEVEL_, weighted by individual parameters β_Effort_ and β_Reward_, reflecting effort and reward sensitivity, respectively (see below for more details). To account for potential time-dependent effects on acceptance rates across repeated trials, the model also incorporated a β_Time_ parameter, which linearly modulated the cost component as a function of trial number (Le Heron et al., 2018). This parameter thus captured time-on-task effects on acceptance rates, whether reflecting fatigue (progressive decline in acceptance across trials) or habituation and rising motivation (progressive increase). The intercept β₀ captured baseline choice bias, reflecting the propensity to accept an offer with zero reward.

After model fitting, we extracted individual β_Effort_ and β_Reward_ parameters as proxies for effort and reward sensitivity, respectively. Parameter estimation was performed using constrained nonlinear optimization (fmincon function in MATLAB 2022b, MathWorks) to minimize the negative log-likelihood of observed choices. Model-derived choice probabilities (range: 0 – 1) were compared against actual behavior.

In this framework, more negative β_Effort_ values indicated stronger effort sensitivity — *i.e.*, a steeper relative decline in acceptance rates with increasing effort. Of note, due to the quadratic nature of the cost function in the winning model (*i.e.*, β_Effort_ × EFFORT_LEVEL_²), this effect is particularly pronounced at high effort levels: the more a participant’s acceptance rate drops for high efforts (*e.g.*, 80% MVC), the steeper the curvature of the effort-sensitivity function and the more negative the β_Effort_. For ease of interpretation in subsequent analyses, β_Effort_ values were multiplied by −1, such that higher (more positive) values consistently reflected stronger effort sensitivity. Conversely, more positive β_Reward_ values corresponded to stronger reward sensitivity, as reflected by steeper relative increase in acceptance with higher rewards. β_Reward_ values were transformed using a log function to normalize the distribution, in accordance with former studies (Le Heron et al., 2018; Derosiere et al., 2024). Subject-specific β_Effort_ and β_Reward_ values were used as dependent variables in voxel-wise, cluster-based permutation analyses (see Statistical Analysis section).

### MRI data analyses

Diffusion data were preprocessed using the Elikopy pipeline (Dessain et al., 2024), including brain extraction (Hoopes et al., 2022), thermal noise removal (Veraart et al., 2016), and corrections for susceptibility-induced distortions, eddy currents, and head motion using FSL (v6.0.7.8). As reversed phase-encoding b0 images were not acquired, susceptibility distortion correction was performed using Synb0-DISCO, which generates a synthetic distortion-free b0 image from the T1-weighted anatomical scan. Post-processing was also performed using Elikopy, which included the mathematical reconstruction of the diffusion-weighted images (64 directions) to derive the diffusion tensor imaging (DTI) model. From this, volume-weighted microstructural maps of FA and MD were generated for each participant in MNI space (Figure 1.C), providing a voxel-wise measure of microstructure across the entire brain.

#### Statistical analyses

As mentioned in the *Introduction*, we used a two-stage strategy to first determine white matter circuits underlying inter-individual differences in effort and reward sensitivity and to then test whether these sensitivities could be decoded from microstructure metrics. First, four whole-brain cluster-based analyses were performed to detect clusters where FA or MD covaried with β_Effort_ or β_Reward_. Second, the FA and MD values from these clusters were used as features in supervised machine learning classifiers to assess whether they could decode individual differences in sensitivity in out-of-sample predictions. These two stages are presented in detail below.

### Whole-brain cluster-based analysis

Voxel-based correlation analyses were conducted using BrainVoyager™ (Brain Innovation, version 23.2), examining the associations between FA and MD maps and β_Effort_ or β_Reward_ (Figure 1.C). These analyses controlled for potential confounding factors by including them as covariates, specifically gender, age, intracranial volume, and other variables that could influence effort and reward sensitivity, such as self-reported depression and anhedonia (assessed using the EDAS and SHAPS scales; (Bonnelle et al., 2016)). Statistical significance was set at a voxel-wise p-value of .005, and correction for multiple comparisons was performed using cluster-size thresholding (threshold: 44 mm^3^; Forman et al., 1995). This approach revealed clusters where white matter microstructure significantly covaried with β_Effort_ or β_Reward_.

To interpret these findings, we use the term white matter *integrity* to describe FA and MD values in our healthy cohort. This term reflects microstructural organization and does not imply pathology. FA and MD provide complementary information: higher FA indicates greater directional coherence of water diffusion, often linked to axonal alignment, density, or myelination, whereas higher MD reflects greater overall diffusivity, potentially due to increased extracellular space or reduced barriers. Because associations may arise in one metric, the other, or both (Hayakawa et al., 2014) analyzing both metrics offers a more comprehensive view of white matter organization. Effects observed across both FA and MD (*i.e.*, with FA and MD clusters overlapping), when replicating in direction, provide stronger evidence for a robust microstructural-behavioral relationship. To align their interpretability, we computed the additive inverse of MD (-MD), allowing FA and -MD to be interpreted in the same direction (higher values indicating greater integrity). For visualization, clusters showing significant negative associations between integrity (FA or -MD) and β_Effort_ or β_Reward_ are highlighted in red, and positive associations in green.

To assign tract labels, we used the XTRACT HCP Probabilistic Tract Atlas (Warrington et al., 2020), which reliably identifies the major canonical white matter bundles, including deep, long-range pathways such as the cingulum, corticospinal tract, and corpus callosum. These tracts, typically located deep, form the principal structural connections within and between hemispheres and are thus the primary focus of our study. A minority of clusters did not overlap with these canonical bundles, instead lying within more superficial white matter (i.e., closer to the cortex) that supports short-range connections between adjacent cortical areas. Because such clusters fall outside the coverage of canonical atlases like XTRACT, we used the EBRAINS Human Connectome Project superficial white matter atlas (Labra-Avila et al., 2020) to facilitate their labelling and describe them in the Supplementary Materials (Supplementary Figures 2 and 3). All tract visualizations presented in this study were derived from white matter bundle definitions and population-averaged templates in MNI ICBM 2009a space, based on previously published work (Garyfallidis et al., 2018; Yeh et al., 2018).

Finally, to characterize the relationship between white matter microstructure and individual differences in effort and reward sensitivity, we conducted partial Pearson correlation analyses. For each significant cluster identified in the voxel-wise analyses, we extracted the FA and -MD values from all voxels within the cluster and computed a mean value per cluster for each participant. These averaged microstructural metrics served as independent variables in separate partial correlation analyses with β_Effort_ and β_Reward_, controlling for the potential confounding factors mentioned above, including age, gender, intra-cranial volume and self-reported measures of depression and anhedonia. The resulting correlation R coefficients, p-values, and corresponding regression plots are presented in the Results section for each cluster. In these plots, the y-axis depicts residualized β_Effort_ or β_Reward_ values (after controlling for covariates), while the x-axis represents the mean FA or -MD values for each cluster, allowing for direct visualization of the strength and direction of the adjusted associations. As before, significant negative and positive partial correlations are shown in red and green, respectively, to facilitate visualization throughout the Results section.

### Decoding effort and reward sensitivity using machine learning

As a second stage of our approach, we used the clusters to train supervised machine learning models, testing whether β_Effort_ and β_Reward_ could be decoded from their microstructural properties. Such predictive analyses are critical, as they provide a principled framework to estimate how focal white matter alterations (e.g., following stroke, surgery, or stimulation) might affect sensitivity to effort and reward, and ultimately disrupt goal-directed behavior. Here, we performed two supervised machine learning analyses, one for β_Effort_ and one for β_Reward_. In each case, we used averaged FA and -MD values of each voxel extracted from the significant clusters identified within major, canonical bundles in the voxel-wise FA and -MD analyses as input features.

Participants were assigned to either a low or high sensitivity class based on a median split of β_Effort_ or β_Reward_ values, enabling binary classification. To avoid bias associated with choosing a single classifier and to ensure robust performance across data distributions, we conducted an algorithmic classifier selection. Specifically, we evaluated 12 classifiers: logistic regression, support vector machines, random forests, gradient boosting, extra trees, adaptive boosting, decision trees, multilayer perceptron, k-nearest neighbors, Gaussian naive Bayes, linear discriminant analysis, and quadratic discriminant analysis.

To estimate classifier performance while avoiding overfitting, we used a nested cross-validation procedure. The outer loop consisted of a stratified 5-fold cross-validation that provided an unbiased estimate of generalization performance. Within each outer fold, a second, inner stratified cross-validation was performed on the training data to conduct a grid search over hyperparameters. The best model identified in the inner loop was then evaluated on the outer test fold. This procedure was repeated 1,000 times, with random re-sampling at each iteration. For each iteration, we computed two key performance metrics: classification accuracy and the area under the curve (AUC) of the receiver operating characteristic curve (ROC). Classification accuracy reflects the proportion of correctly predicted labels, while the ROC plots the true positive rate against the false positive rate across all decision thresholds. The AUC gives a synthetic and quantitative measure of this curve, with 0.5 indicating chance-level performance and 1.0 representing perfect classification.

For each target variable (β_Effort_ and β_Reward_), the classifier achieving the highest mean AUC across the 1,000 iterations was selected for further analysis. The best-performing classifier was extra trees for β_Effort_ and logistic regression for β_Reward_, although all classifiers performed above chance as described in the Results section.

To assess whether our classification accuracy and AUC was significantly above chance level, we repeated the entire classification procedure using the best-performing classifiers 1,000 times with randomly permuted class labels, generating a null distribution of accuracy and AUC under the assumption of no association between white matter microstructure and β_Effort_ and β_Reward_. We then evaluated whether classification accuracy and AUC in the true-label data exceeded that expected by chance using three complementary criteria: (1) a Monte Carlo p-value, defined as the proportion of classification yielding higher accuracy or AUC with the permuted-label data than that obtained with the true-label data; (2) one-sided t-tests comparing the distributions of accuracy or AUC for true-label versus permuted-label data; and (3) one-sided t-tests testing whether accuracy or AUC for true-label and permuted-label data were significantly higher from the chance level of 0.5. Statistical significance was set at p < .05.

Further, to determine whether the observed classification performance was specific to the white matter clusters identified in the voxel-wise analysis, we conducted a control analysis using randomly selected clusters. For each of the two target variables (β_Effort_ and β_Reward_), we randomly sampled clusters from the whole-brain white matter mask, matching both the number and size of the original significant clusters. We then repeated the full classification pipeline, including evaluation across 12 classifiers, nested cross-validation, and performance estimation based on accuracy and AUC, for these randomly located clusters. As with the primary analyses, we assessed whether AUC exceeded chance (0.5) using one-sample t-tests; p-values were FDR-corrected. This control analysis allowed us to test whether decoding performance was specific to the significant clusters identified in the voxel-wise analyses, rather than reflecting generic variability of white matter microstructure across subjects.

Finally, we conducted feature importance and recursive feature elimination analyses to identify which clusters contributed most to classification performance. In this context, the term “feature” refers specifically to the individual white matter clusters included as input variables in the model, a standard term in machine learning used to denote each parameter or predictor considered during the analysis. For feature importance, we used the best-performing classifier for each target variable (extra trees for β_Effort_, logistic regression for β_Reward_) trained on true-label data. We extracted built-in feature weights (i.e., impurity-based importance scores for tree-based models and absolute coefficient weights for logistic regression) to rank all identified clusters’ microstructural measures (FA and -MD) according to their predictive contribution. Using these rankings, we implemented recursive feature elimination by iteratively removing the least informative cluster and repeating the full classification analysis, recalculating AUC at each iteration. This process was repeated 100 times with random re-sampling to ensure stability of the performance estimates. For each number of retained clusters (from one up to the full set), we computed the mean AUC and tested whether AUC significantly exceeded chance level (0.5) using one-sided t-tests across iterations. This analysis provided a quantitative estimate of the optimal subset of clusters required to decode β_Effort_ and β_Reward_ above chance, while identifying the specific clusters driving classification performance.

## RESULTS

### Effort and reward sensitivity show substantial inter-individual variability across healthy subjects

Behavioral data revealed strong inter-individual differences in how participants modulated their acceptance rates based on effort and reward magnitudes, reflecting differences in effort and reward sensitivity (Figure 2.A). This variability was quantitatively captured by the density distribution of the β_Effort_ and β_Reward_ parameters, which showed broad dispersion across individuals (Figure 2.B; β_Effort_: mean = 16.92, SD = 11.10; β_Reward_: mean = -0.30, SD = 0.98). The coefficient of variation reached 65.6% for β_Effort_ and 318.1% for β_Reward_, confirming substantial heterogeneity in sensitivity to effort and reward. To illustrate this diversity of profiles, Figure 2.C presents individual examples of acceptance behavior in participants with low vs. high β_Effort_ and β_Reward_ values. These findings confirm the presence of heterogenous sensitivity profiles in the healthy population.

**Figure 2:**
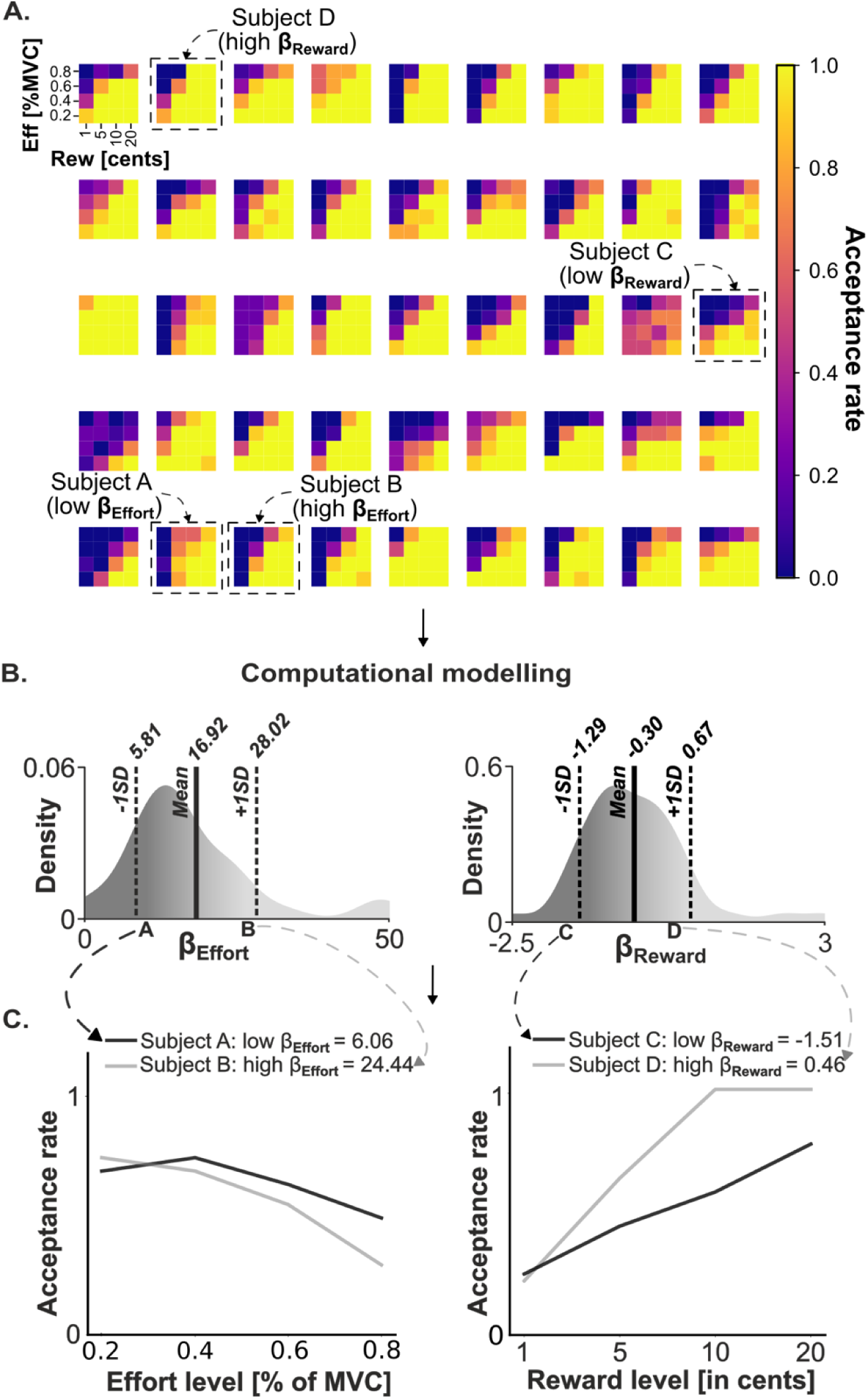
Effort and reward sensitivity show substantial inter-individual variability across healthy subjects. **A. Individual acceptance rate maps.** Each map represents the acceptance rate of a given subject as a function of effort (in % MVC; y-axes) and reward (x-axes). These maps show the strong inter-individual differences in how participants modulated their acceptance rates based on effort and reward magnitudes. **B. Distributions of β_Effort_ and β_Reward_.** Parameters β_Effort_ and β_Reward_ were estimated from the modelling of acceptance rates presented in A using the equation presented in section *Computational modelling of acceptance rates*. Here again, the density distributions depict the high inter-individual variability in β_Effort_ and β_Reward_ values among our group of 45 subjects. As described in the section *Computational modelling of acceptance rates*, for ease of interpretation, β_Effort_ values initially obtained following our modelling procedure were multiplied by −1, such that higher (more positive) values reflect stronger effort sensitivity in the left panel. Further, more positive β_Reward_ values correspond to stronger reward sensitivity in the right panel. **C. Representative acceptance curves from individuals with divergent β_Effort_ and β_Reward_ values**. *Left panel:* Compared to subject A, subject B showed a stronger decrease in acceptance rates with increasing effort, reflected by a higher β_Effort_ (24.44 vs. 6.06). *Right panel*: Compared to subject C, subject D showed a stronger increase in acceptance rates with increasing reward, reflected by a higher β_Reward_ (0.46 vs. –1.51). These examples illustrate how modulation of decision behavior by effort and reward varies across healthy individuals.

### Key findings on the microstructural correlates of effort and reward sensitivity

As detailed in the *Methods*, in the first stage of our statistical approach, we conducted four cluster-based analyses to examine the microstructural correlates of individual differences in β_Effort_ and β_Reward_, using FA and the additive inverse of MD (- MD, with higher values reflecting greater integrity) to capture complementary aspects of white matter integrity (see section *Whole-brain cluster-based analysis*, for more details). Across these analyses, we identified 12 significant clusters located within major, canonical white matter bundles: 5 covarying with effort sensitivity and 7 with reward sensitivity. All of the 5 effort-related clusters were situated within tracts connected to frontal valuation regions, such as SMA, dACC, and OFC (*e.g*., within the anterior cingulum tract). In contrast, the 7 reward-related clusters were more widely distributed: 3 were located within tracts linked to frontal valuation regions, while 4 lay outside this network, encompassing fronto-parietal and sensorimotor pathways. The most robust associations included one cluster shared across effort and reward sensitivity analyses and another consistent across FA and -MD metrics, both within SMA-connected tracts. Machine learning analyses further showed that microstructural metrics from these 5 effort-related and 7 reward-related clusters reliably predicted effort and reward sensitivity, respectively, whereas randomly positioned clusters did not. Table 1 summarizes all clusters identified across the four analyses (i.e., FA and - MD analyses for β_Effort_ and β_Reward_), their tract labels and associated partial correlation results, which are described individually in the following sections.

**Table 1.** White matter clusters showing significant associations with β_Effort_ and β_Reward_. The table reports 12 significant clusters (5 associated with β_Effort_ and 7 with β_Reward_). For each cluster, MNI coordinates correspond to the cluster center, and volume reflects cluster size (mm^3^). Tract labels were derived using the XTRACT HCP Probabilistic Tract Atlas and, when necessary, refined with anatomical masks from prior studies. The two rightmost columns report partial correlation coefficients (R) and p-values between β_Effort_ or β_Reward_ and the average FA or -MD values extracted from voxels within each cluster, controlling for age, gender, and other covariates (see Methods). Clusters located within tracts directly connected to frontal valuation regions (*e.g.*, projecting to the SMA or OFC) are highlighted in black, while clusters belonging to fronto-parietal or sensorimotor pathways are shown in gray.

| MNI coordinates |  |  | Volume<br>(mm<sup>3</sup>) | Tract label | Partial correlations |  |
| --- | --- | --- | --- | --- | --- | --- |
| x | y | z |  |  | R value | p-value |
| <u>5 clusters showing significant associations between white matter integrity and $\beta_{\text{Effort}}$</u> |  |  |  |  |  |  |
| FA analysis |  |  |  |  |  |  |
| -13.95 | -10.01 | 58.02 | 94 | SMA portion of the left corticospinal tract | -0.697 | < .001 |
| -10.93 | 15.96 | 19.02 | 56 | Left mid-anterior segment of the corpus callosum | -0.556 | < .001 |
| -9.95 | 34.7 | 11.22 | 209 | Left anterior cingulum bundle | 0.623 | < .001 |
| -MD analysis |  |  |  |  |  |  |
| -7.55 | 14.03 | 20.11 | 322 | Left mid-anterior segment of the corpus callosum | -0.641 | < .001 |
| 8.71 | 15.55 | 20.32 | 139 | Right mid-anterior segment of the corpus callosum | -0.575 | < .001 |
| <u>7 clusters showing significant associations between white matter integrity and $\beta_{\text{Reward}}$</u> |  |  |  |  |  |  |
| FA analysis |  |  |  |  |  |  |
| -13.81 | -10.44 | 59.95 | 79 | SMA portion of the left corticospinal tract | -0.614 | < .001 |
| -17.82 | 53.12 | -2.3 | 73 | Left forceps minor | 0.561 | < .001 |
| 17.9 | 53.81 | 4.19 | 67 | Right forceps minor | 0.572 | < .001 |
| 39.85 | 15.76 | 17.7 | 126 | Right superior lateral fasciculus | 0.579 | < .001 |
| 36.22 | -24.6 | 33.78 | 86 | Right superior lateral fasciculus | 0.561 | < .001 |
| -9.61 | 5.02 | 26.19 | 137 | Left corpus callosum (motor portion) | 0.566 | < .001 |
| -MD analysis |  |  |  |  |  |  |
| 11.05 | -29.94 | -34.32 | 84 | Right middle cerebellar peduncle /<br>left cortico-ponto-cerebellum tract | -0.547 | < .001 |

### Microstructure in tracts connected to key frontal valuation regions is associated with effort sensitivity

Among the 5 significant clusters associated with β_Effort_, 4 showed negative associations, where reduced FA or -MD (*i.e.*, lower microstructural integrity) was linked to higher β_Effort_ (*i.e.*, stronger effort sensitivity), while 1 showed a positive association. Below, we detail these clusters.

The first negative association with β_Effort_ was found for FA in a cluster located within the SMA portion of the left corticospinal tract (MNI: -13.95, -10.01, 58.02; 94 mm_3_; R = -.697; p < .001; Figure 3A). Probabilistic atlas labelling identified the corticospinal tract, and anatomical masks (Bonnelle et al., 2016; Beckmann et al., 2009) showed it was located within SMA proper. As typical in such labelling, the cluster extended beyond the tract (46% of voxels in the tract), likely involving other SMA-related projections, including the SMA-NAcc pathway, which showed in prior work a negative association between effort sensitivity and macroscopic structural connectivity (i.e., streamline count; Derosiere et al., 2024). Importantly, the same cluster also emerged as a key correlate of β_Reward_ (see below), underscoring its importance as a shared microstructural substrate of both effort and reward sensitivity.

**Figure 3:**
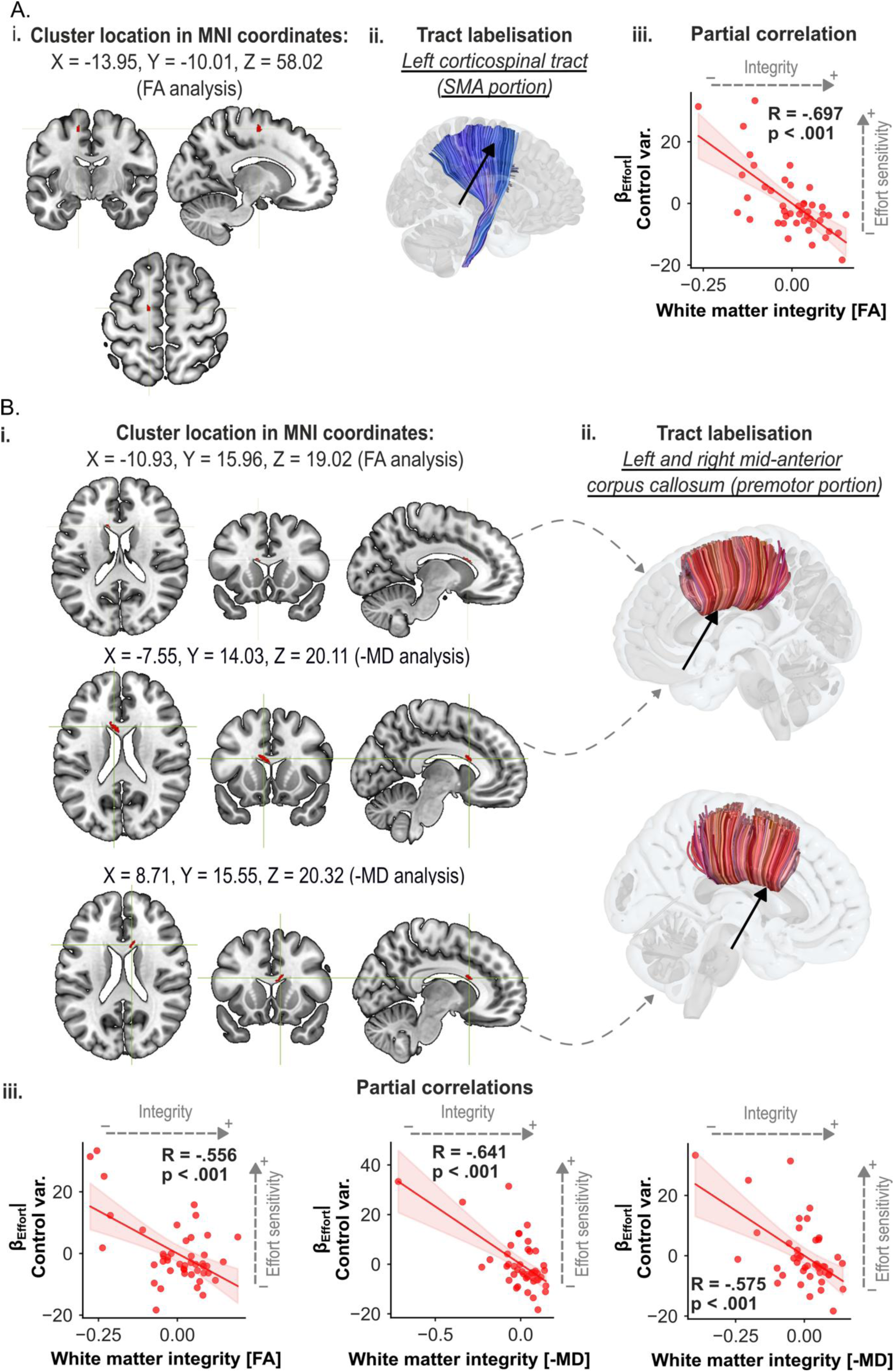
Microstructure in tracts connected to frontal valuation regions showing negative associations with effort sensitivity (β_Effort_). **A. Cluster in the SMA portion of the left corticospinal tract.** (i) A significant cluster (red) showed a negative correlation between FA and β_Effort_ (MNI: -13.95, -10.01, 58.02). (ii) Probabilistic tract labelling identified the corticospinal tract; anatomical masks (Bonnelle et al., 2016; Beckmann et al., 2009) confirmed the cluster’s location within SMA proper. The cluster extended beyond the tract (46% of its voxels), likely encompassing SMA-related projections such as the SMA–NAcc pathway, previously linked to β_Effort_ via macroscopic connectivity measures (Derosiere et al., 2024). (iii) Partial correlation confirmed reduced FA in this cluster was significantly correlated with higher effort sensitivity (R = -.697, p < .001). Across all panels, residualized β values (adjusted for age, gender, intracranial volume, depression, and anhedonia) are plotted against mean FA or -MD per cluster to visualize adjusted associations. **B. Clusters in the mid-anterior corpus callosum.** (i) Three significant clusters (red) exhibited consistent negative correlations between microstructural integrity and β_Effort_. The FA cluster (top row, left) overlapped with one -MD cluster (middle row, left), while another -MD cluster was located symmetrically in the right hemisphere (bottom row). (ii) Probabilistic tract labelling localized all clusters to the mid-anterior corpus callosum, specifically in its premotor portion containing interhemispheric fibers connecting bilateral SMAs (Xiong et al., 2024). (iii) Partial correlation analyses confirmed strong negative associations for each cluster (R = -.556, R = -.641, and R = -.575, all p < .001). The replication across hemispheres and both FA and -MD provides particularly strong evidence for a robust link between effort sensitivity and compromised integrity in SMA-related interhemispheric pathways.

A second robust pattern involved the mid-anterior corpus callosum. FA analyses revealed a cluster in its left portion (MNI: -10.93, 15.96, 19.02; 56 mm_3_; R = -.556; p < .001; Figure 3B), in a locus containing interhemispheric fibers predominantly connecting bilateral SMAs (Xiong et al., 2024). Supporting this, -MD analyses revealed two additional bilateral clusters with overlapping spatial distributions with the FA cluster (left: -7.55, 14.03, 20.11; 322 mm_3_; R = -.641; p < .001; right: 8.71, 15.55, 20.32; 139 mm_3_; R = -.575; p < .001; Figure 3B). The fact that these effects replicate bilaterally and across both FA and -MD, consistently showing decreased integrity with higher β_Effort_, provides particularly strong evidence for a robust relationship. These results suggest that higher effort sensitivity is linked to reduced directional integrity (lower FA, reflecting reduced axonal alignment or myelination) and increased water diffusion (lower -MD, suggesting greater extracellular space), which may disrupt interhemispheric communication between bilateral SMAs and other medial frontal regions critical for effort processing.

In contrast, one cluster showed a positive association between FA and β_Effort_. This cluster was located in the left anterior cingulum bundle (MNI: -9.95, 34.7, 11.22; 209 mm_3_; R = .623; p < .001; Figure 4), a tract that links medial frontal regions. To identify which medial frontal regions were specifically concerned by this cluster, we projected it onto a high-resolution atlas (Labra-Avila et al., 2020), which revealed it was located in fibers connecting the dACC with SMA and medial OFC, key frontal valuation hubs. No additional clusters emerged in major tracts, and the -MD analysis revealed none with positive associations.

**Figure 4:**
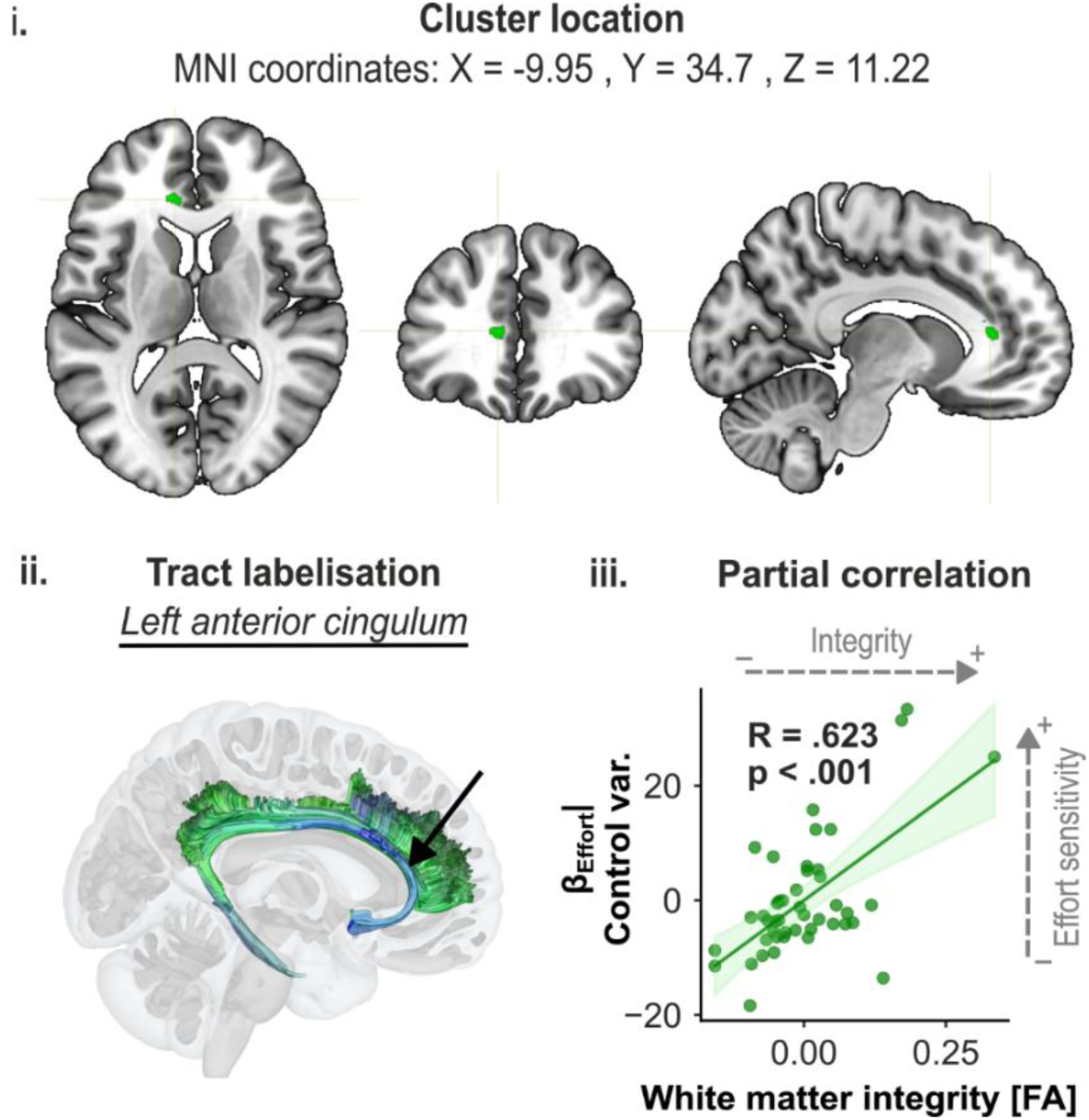
Microstructure in a tract connected to frontal valuation regions showing a positive association with effort sensitivity (β_Effort_). (i) A significant cluster (green) exhibited a positive correlation between FA and β_Effort_ (MNI: -9.95, 34.7, 11.22). (ii) Probabilistic tract labelling identified the anterior cingulum. To clarify which medial frontal regions were specifically connected by this cluster, we projected it onto a high-resolution atlas (Labra-Avila et al., 2020), revealing fibers linking the dACC with SMA and medial OFC, core valuation hubs. (iii) Partial correlation confirmed that greater FA in this cluster was associated with higher effort sensitivity (R = .623, p < .001). No additional clusters were identified in major tracts, and the - MD analysis did not reveal any clusters with positive associations.

Together, these findings indicate that individual differences in effort sensitivity are anchored in several white matter pathways connected to frontal valuation regions. Reduced microstructural integrity in SMA-connected clusters, including clusters identified in the corticospinal tract and interhemispheric fibers of the mid-anterior corpus callosum, is consistently associated with stronger effort sensitivity, representing some of the most robust effects in the present study (consistent across metrics and, for one cluster, across valuation dimensions [see below]). Conversely, increased integrity of the left anterior cingulum bundle, encompassing connections between SMA, dACC, and medial OFC, is associated heightened effort sensitivity, potentially reflecting a distinct mechanism whereby enhanced communication among these valuation hubs amplifies effort sensitivity (see *Discussion* section).

### Microstructure in tracts connected to frontal valuation regions, as well as fronto- parietal and sensorimotor structures, is associated with reward sensitivity

As mentioned earlier, the FA and -MD analyses revealed 7 significant clusters, more widely distributed than the effort-related clusters: 3 were located within tracts linked to frontal valuation regions, while 4 lay outside this network, encompassing fronto-parietal and sensorimotor pathways. 2 clusters showed negative associations, where lower FA or -MD (*i.e.*, lower microstructural integrity) was linked to higher β_Reward_ (*i.e.*, stronger reward sensitivity), while 5 clusters showed positive associations. Below, we detail these clusters, beginning again with the negative associations.

The first negative association with β_Reward_ was for FA in a cluster located within the SMA portion of the left corticospinal tract (MNI: -13.81, -10.44, 59.95; 79 mm_3_; R = -.614; p < .001; Figure 5A), as identified with probabilistic atlas labelling and anatomical masks (Bonnelle et al., 2016; Beckmann et al., 2009) and overlapped extensively with the cluster identified for β_Effort_ (MNI: -13.95, -10.01, 58.02; 94 mm_3_; Figure 6). Importantly, this shared cluster represents the sole locus where microstructural integrity covaries with both β_Effort_ and β_Reward_, suggesting a shared white matter substrate where reduced integrity may amplify both effort and reward sensitivity. To confirm that this dual association was not trivially driven by the fact that β_Effort_ and β_Reward_ were derived from the same acceptance rates, we tested whether FA values in these clusters correlated with the two other model-derived parameters: β_Time_ and β₀. No significant correlations were observed either with the effort-related FA values (β_Time_: R = -.180, p = .237; β₀: R = -.163, p = .284) or with the reward-related FA values (β_Time_: R = -.111, p = .466; β₀: R = -.148, p = .332; Supplementary Figure 4), underscoring the specificity of this SMA white matter locus for β_Effort_ and β_Reward_.

**Figure 5:**
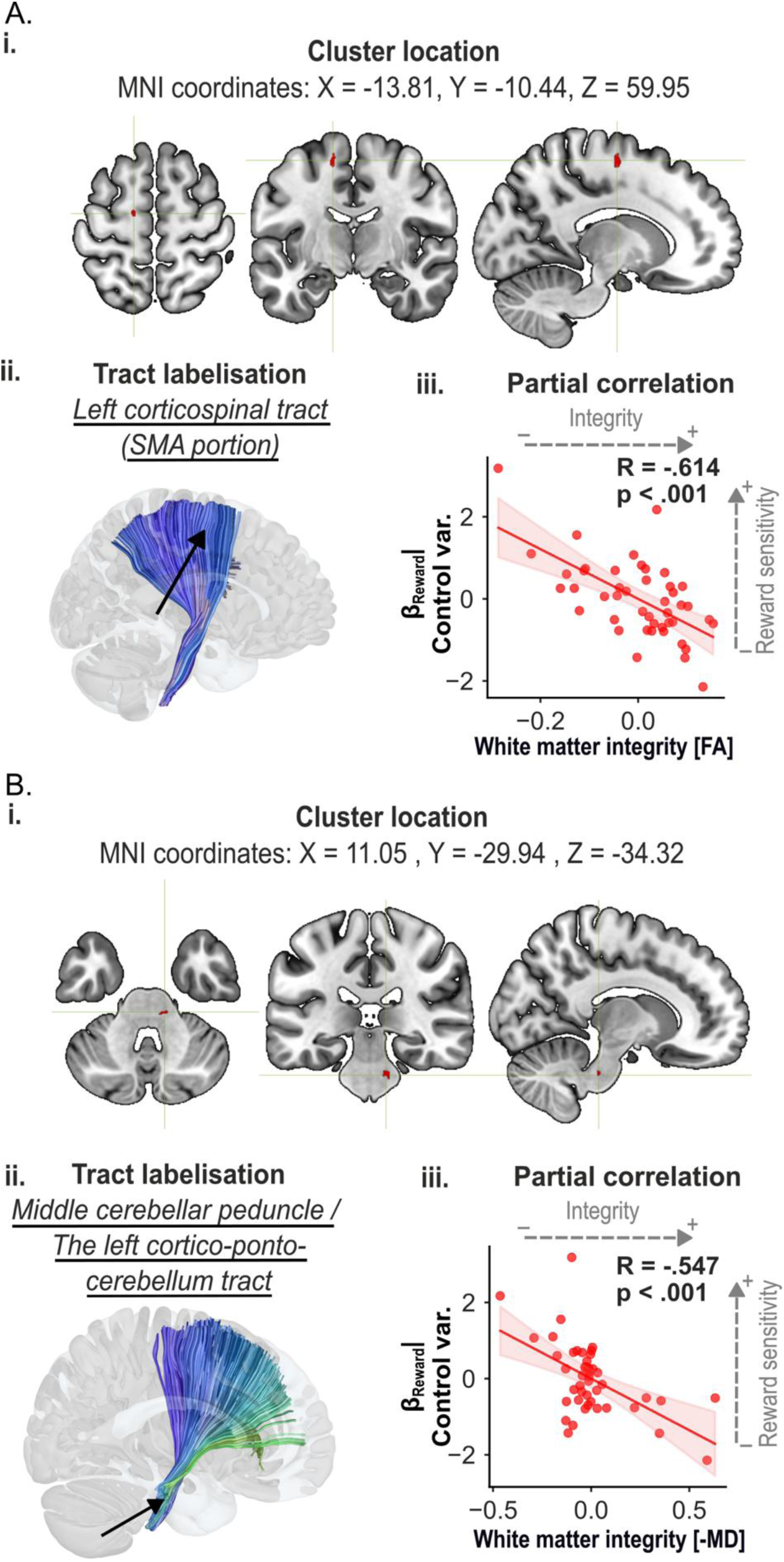
Microstructure in tracts connected to frontal valuation regions and sensorimotor structures and showing negative associations with reward sensitivity (β_Reward_). **A. Cluster in the SMA portion of the left corticospinal tract.** (i) A significant cluster (red) exhibited a negative correlation between FA and β_Reward_ (MNI: -13.81, -10.44, 59.95). (ii) Probabilistic tract labelling identified the corticospinal tract; anatomical masks (Bonnelle et al., 2016; Beckmann et al., 2009) confirmed the cluster’s location within SMA proper. The cluster overlapped substantially with the β_Effort_ cluster (see Figure 6), suggesting a shared white matter substrate where reduced integrity may amplify both effort and reward sensitivity. (iii) Partial correlation confirmed that reduced FA in this region was significantly correlated with greater reward sensitivity (R = -.614, p < .001). **B. Cluster in the right middle cerebellar peduncle, extending to the left cerebello-ponto-cortical tract.** (i) A significant cluster (red) exhibited a negative correlation between -MD and β_Reward_ (MNI: 11.05, -29.94, -34.32). (ii) Probabilistic tract labelling localized the cluster right middle cerebellar peduncle, extending to the left cerebello-ponto-cortical tract. (iii) Partial correlation confirmed that reduced integrity in this region was significantly correlated with greater reward sensitivity (R = -.547, p < .001). Across all panels, residualized β values (adjusted for age, gender, intracranial volume, depression, and anhedonia) are plotted against mean FA or -MD per cluster to visualize adjusted associations.

**Figure 6.**
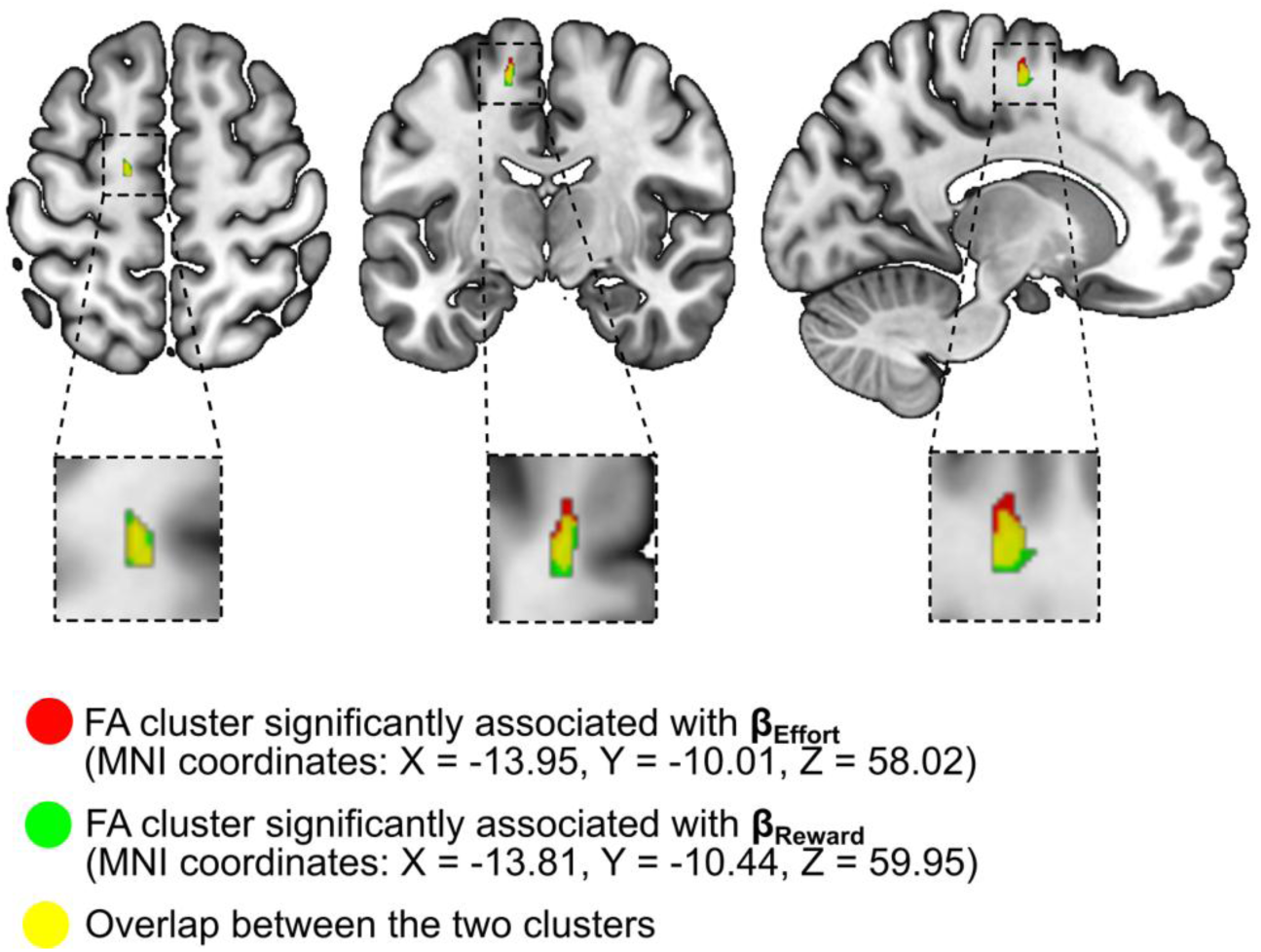
Exclusive overlap of white matter clusters associated with effort and reward sensitivity within the SMA portion of the corticospinal tract. FA analyses identified two significant clusters within the SMA portion of the left corticospinal tract. The cluster associated with β_Effort_ (red) was centered at MNI: X = –13.95, Y = –10.01, Z = 58.02 (94 mm_3_), while the cluster associated with β_Reward_ (green) was centered at MNI: X = –13.81, Y = –10.44, Z = 59.95. These clusters overlapped extensively (yellow), forming the only locus across the entire FA and -MD maps where microstructural integrity covaried with both β_Effort_ and β_Reward_. This convergence underscores the specificity of this SMA corticospinal segment as a shared structural substrate for individual differences in sensitivity to both effort and reward.

The second negative association was located in a tract connected to sensorimotor structures, specifically the right middle cerebellar peduncle, with part of the cluster extending into the left cortico-ponto-cerebellar tract, which projects to the middle cerebellar peduncle, as identified by probabilistic tract labelling (MNI: 11.05, –29.94, –34.32; 84 mm_3_; R = –.547; p < .001; Figure 5B). Here, lower -MD values, indicative of greater extracellular space, was associated with higher β_Reward_. Although less expected, this result suggests that disrupted cortical input to the cerebellum (*i.e.*, since the middle cerebellar peduncle conveys afferences from cortex to cerebellum) may promote action initiation when high rewards are at stakeThis finding illustrates how a whole-brain, cluster-based approach applied to FA and -MD maps can uncover functionally relevant pathways beyond those typically targeted by tract-of-interest analyses.

5 clusters showed positive associations between FA and β_Reward_. Two bilateral clusters were located in the anterior portion of the forceps minor (left: MNI: -17.82, 53.12, -2.3; 73 mm_3_; R = .572; p < .001; right: 17.9, 53.81, 4.19; 67 mm_3_; R = .561; p < .001; Figure 7A), a tract interconnecting bilateral OFCs (Filbey et al., 2014), key hubs for reward valuation. Two additional clusters were found in a fronto-parietal pathway, namely along the right superior longitudinal fasciculus, one in its frontal portion (MNI: 39.85, 15.76, 17.7; 126 mm_3_; R = .579; p < .001) and one in its parietal portion (MNI: 36.22, -24.6, 33.78; 86 mm_3_; R = .561; p < .001; Figure 7B). Finally, one other cluster was located in a tract connected to motor structures, namely in the left mid-body of the corpus callosum (MNI: -9.61, 5.02, 26.19; 137 mm_3_; R = .566; p < .001; Figure 7C), a locus of fibers primarily connecting bilateral M1s (Hofer & Frahm, 2006; Tarumi et al., 2022; Wahl et al., 2007). No additional clusters emerged in major tracts, and the -MD analysis revealed none with positive associations.

**Figure 7:**
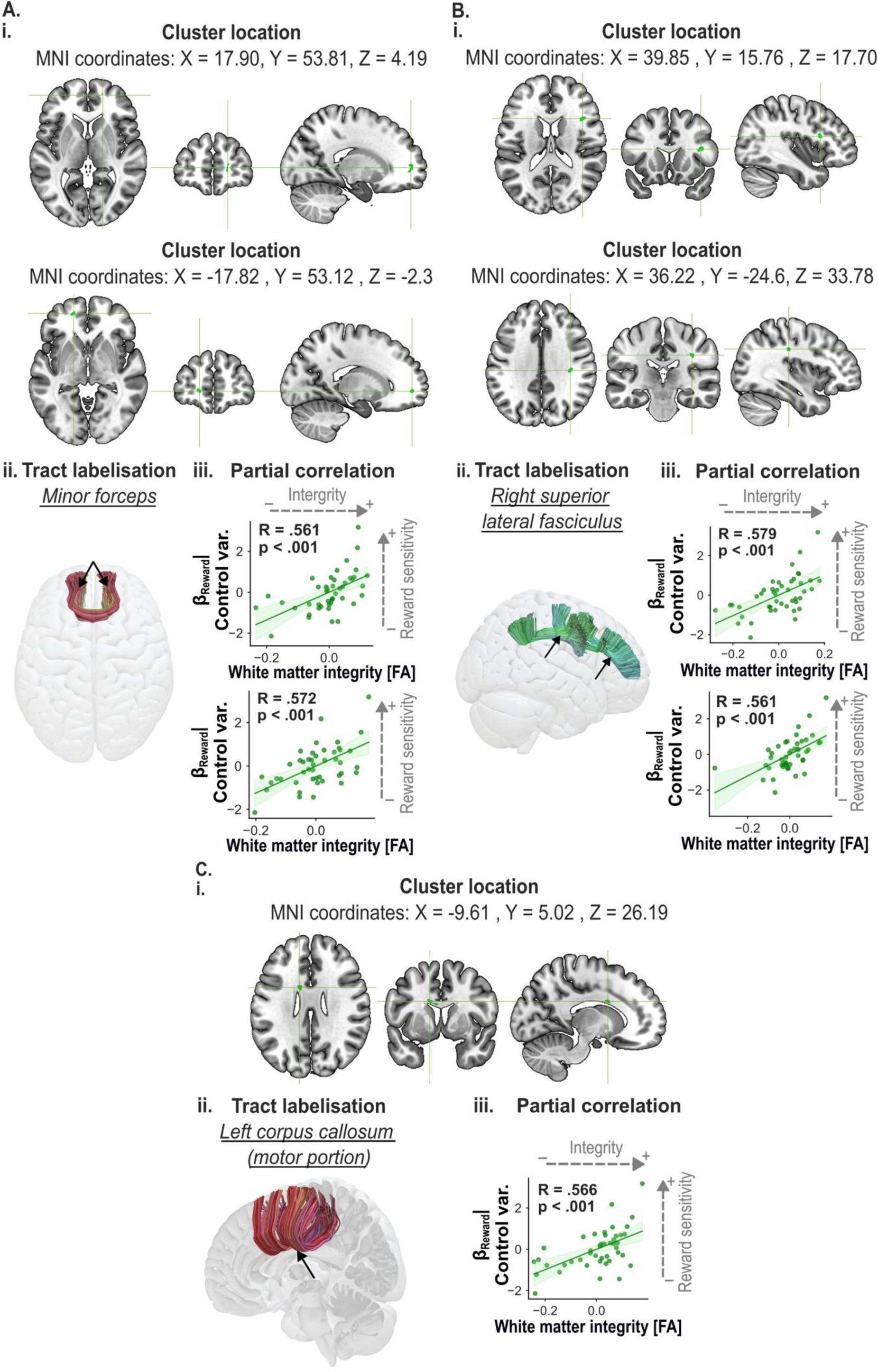
Microstructure in tracts connected to frontal valuation regions, fronto-parietal and sensorimotor structures showing positive associations with reward sensitivity (β_Reward_). **A. Bilateral clusters in the forceps minor.** (i) Two significant clusters (green) exhibited a positive correlation between FA and β_Reward_ (MNI: left: -17.82, 53.12, -2.3; right: 17.9, 53.81, 4.19). (ii) Probabilistic tract labelling localized both clusters to the forceps minor, interconnecting bilateral OFCs, key hubs for reward valuation. (iii) Partial correlation confirmed significant positive associations with reward sensitivity (R = .572 and R = .561, both p < .001). **B. Clusters in the right superior longitudinal fasciculus.** (i) Two significant clusters (green) exhibited a positive correlation between FA and β_Reward_ (MNI: anterior: 39.85, 15.76, 17.7; posterior: 36.22, -24.6, 33.78). (ii) Probabilistic tract labelling indicated both clusters were located within the superior longitudinal fasciculus. (iii) Partial correlation confirmed significant associations with reward sensitivity (R = .579 and R = .561, both p < .001). **C. Cluster in the left mid-body of the corpus callosum (motor portion).** (i) One significant cluster (green) exhibited a positive correlation between FA and β_Reward_ (MNI: -9.61, 5.02, 26.19). (ii) Probabilistic tract labelling indicated the cluster was located in callosal fibers connecting bilateral motor cortices. (iii) Partial correlation confirmed a significant association (R = .566, p < .001).

Collectively, these results show that differences in reward sensitivity are associated with white matter integrity across distributed systems, encompassing circuits directly connected to frontal valuation regions as well as fronto-parietal and sensorimotor pathways. Regarding frontal valuation regions, heightened sensitivity was linked to reduced integrity within an SMA cluster, comprising SMA-connected corticospinal fibers, and increased integrity within the OFC-connected anterior forceps minor. In addition, reduced integrity in cerebellum-connected tracts and greater integrity within a major fronto-parietal bundle (i.e., the superior longitudinal fasciculus) and motor callosal fibers was linked to stronger reward sensitivity, highlighting additional contributions from fronto-parietal and sensorimotor pathways. This pattern suggests that reward sensitivity reflects the integrated influence of frontal valuation circuits together with fronto-parietal and motor-related networks, rather than any single network.

### Decoding effort and reward sensitivity from brain microstructure using machine learning classifiers

We next tested whether β_Effort_ and β_Reward_ could be predicted from the microstructural properties of the clusters identified above. Our machine learning models used as input the averaged FA and -MD values extracted from the significant clusters identified in voxel-wise analyses and were trained to classify participants into low versus high β_Effort_ and β_Reward_ groups. Performance metrics (accuracy and AUC) were estimated using nested cross-validation on true labels and compared against null distributions generated by 1,000 label permutations.

For β_Effort_, the classifier achieved a mean accuracy of 0.648 ± 0.053 on the true-label data, well above the 0.5 chance level for binary classification. In contrast, accuracy on the permuted-label data averaged 0.506 ± 0.086, with only 37 out of 1,000 permutations reaching or exceeding the accuracy obtained with true-label data, yielding a Monte Carlo p-value of .037 (Figure 8.A). This difference in accuracy between true-label and permuted-label data was statistically significant (t_998_ = 45.57, p < .001, Cohen’s d > 2.5), and accuracy on true-label data was significantly above chance (t_999_ = 88.19, p < .001), unlike the permuted-label data (t_999_ = -1.466, p = .93). To complement these results, we also examined the AUC, which provides a threshold-independent measure of classification performance. The AUC obtained on true-label data was 0.719 ± 0.050, higher than the AUC from permuted-label data (0.498 ± 0.118), with only 28 out of 1,000 permutations matching or exceeding it (Monte Carlo p = .028). The difference between true-label and permuted-label data was also statistically significant (t_998_ = 54.40, p < .001, Cohen’s d > 2.5), and only the AUC obtained on the true-label data was significantly greater than chance (t_999_ = 137.3, p < .001; permuted-label data: t_999_ = -0.553, p = .580).

**Figure 8:**
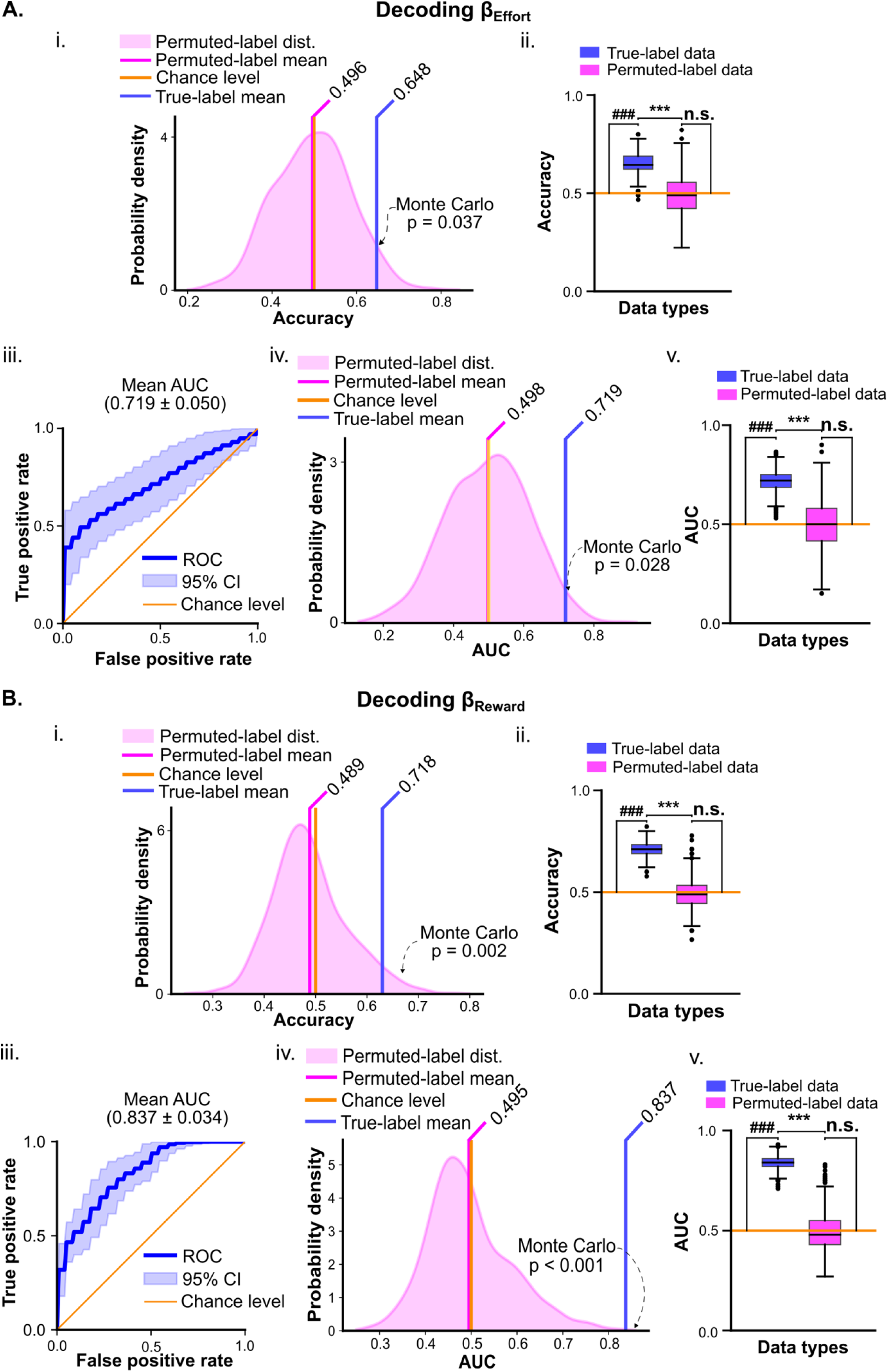
White matter microstructure predicts individual differences in effort and reward sensitivity. **(A) β_Effort_ decoding.** (i) Classifier accuracy on true-label data (blue line) compared to the null distribution obtained from 1,000 label permutations (pink); only 37/1,000 permutations matched or exceeded true-label accuracy (Monte Carlo p = 0.037). (ii) Accuracy was significantly higher for true-label data compared to permuted-label and chance-level data (true vs. permuted: t_998_ = 45.57, P < 0.001; true vs. chance: t_999_ = 88.19, P < 0.001; permuted vs. chance: t_999_ = –1.466, P = 0.93). (iii) ROC curve showing classifier performance (mean AUC = 0.720 ± 0.039; 95% CI shown in shading). (iv) True-label AUC (blue line) compared to null distribution from permuted-label data (pink); 28/1,000 permutations matched or exceeded true-label AUC (Monte Carlo p = 0.028). (v) AUC was significantly higher for true-label data (true vs. permuted: t_998_ = 54.40, P < 0.001; true vs. chance: t_999_ = 137.3, P < 0.001; permuted vs. chance: t_999_ = –0.553, P = 0.580). **(B) β_Reward_ decoding**. (i) True-label accuracy (blue line) exceeded that of 1,000 permutations (pink); only 2 permutations matched or exceeded it (Monte Carlo P = 0.002). (ii) Accuracy was significantly higher for true-label data (true vs. permuted: t_998_ = 88.68, P < 0.001; true vs. chance: t_999_ = 183.2, P < 0.001; permuted vs. chance: t_999_ = –4.95, P = 0.99). (iii) ROC curve (mean AUC = 0.829 ± 0.022). (iv) AUC distribution from permuted-label data (pink) with true-label AUC (blue line); no permutations matched or exceeded the true-label AUC (Monte Carlo P < 0.001). (v) AUC was significantly higher for true-label data (true vs. permuted: t_998_ = 112.7, P < 0.001; true vs. chance: t_999_ = 314.5, P < 0.001; permuted vs. chance: t_999_ = –1.698, p = 0.955). Boxplots show median, interquartile range, and full range. Symbols including # and * indicate statistical significance: ^###^p < .001, ***p < .001; n.s. = not significant)

Similarly, for β_Reward_, the classifier achieved a mean accuracy of 0.718 ± 0.038 on the true-label data. In contrast, accuracy on the permuted-label data averaged 0.489 ± 0.073, with only 2 out of 1,000 permutations reaching or exceeding the accuracy obtained with true-label data, yielding a Monte Carlo p-value of .002 (Figure 8.B). This difference between true-label and permuted-label data was statistically significant (t_998_ = 88.68, p < .001, Cohen’s d > 2.5), and accuracy on true-label data was significantly above chance (t_999_ = 183.2, p < .001), unlike the permuted-label data (t_999_ = -4.95, p = .99). The AUC from true-label data was 0.837 ± 0.033, while the one from permuted data-label was 0.495 ± 0.090, with 0 out of 1,000 permutations matching or exceeding it (Monte Carlo p < .001). This difference was also statistically significant (t_998_ = 112.7, p < .001, Cohen’s d > 2.5), and only the AUC from true-label data was significantly greater than chance (t_999_ = 314.5, p < .001; permuted-label data: t_999_ = -1.698, p = .955).

To further establish the robustness and specificity of our machine learning findings, we conducted two additional analyses. First, we verified that decoding performance was not dependent on the choice of the best classifier by repeating the t-tests against chance level across the AUC values obtained for all 12 tested classifiers (i.e., with p values FDR-corrected). AUC was significantly above chance level across nearly all classifiers for β_Effort_ (all p-values < .001, except SVM: p = .999) and across all classifiers for β_Reward_ (all p-values < .001; see Figure 9 & Supplementary Tables 1-2), underscoring that predictive performance was classifier-independent and generalizable. Second, we tested whether decoding performance was specific to the white matter clusters identified in our voxel-wise analyses. For each target variable, we randomly sampled clusters from the whole-brain white matter mask, matched in number and size to the original clusters, and re-ran the full classification pipeline, including evaluation across all classifiers and nested cross-validation. In contrast to the primary analyses, AUC for these randomly located clusters was consistently near chance (β_Effort_ prediction: AUC values = [.364 to .506], all p-values = [.360 to 1.00]; β_Reward_ prediction: AUC values = [.401 to .534], all p-values = [.368 to 1.00]; Figure 9, pink). In fact, the AUC obtained from the original, significant clusters with the best-performing classifier was significantly higher than the AUC obtained from the randomly located clusters (t_999_ = 192.4, p < .001). This second analysis demonstrates that the decoding of β_Effort_ and β_Reward_ is driven by microstructural features of the specific clusters identified in the voxel-wise analyses, rather than reflecting generic inter-individual variability in white matter.

**Figure 9:**
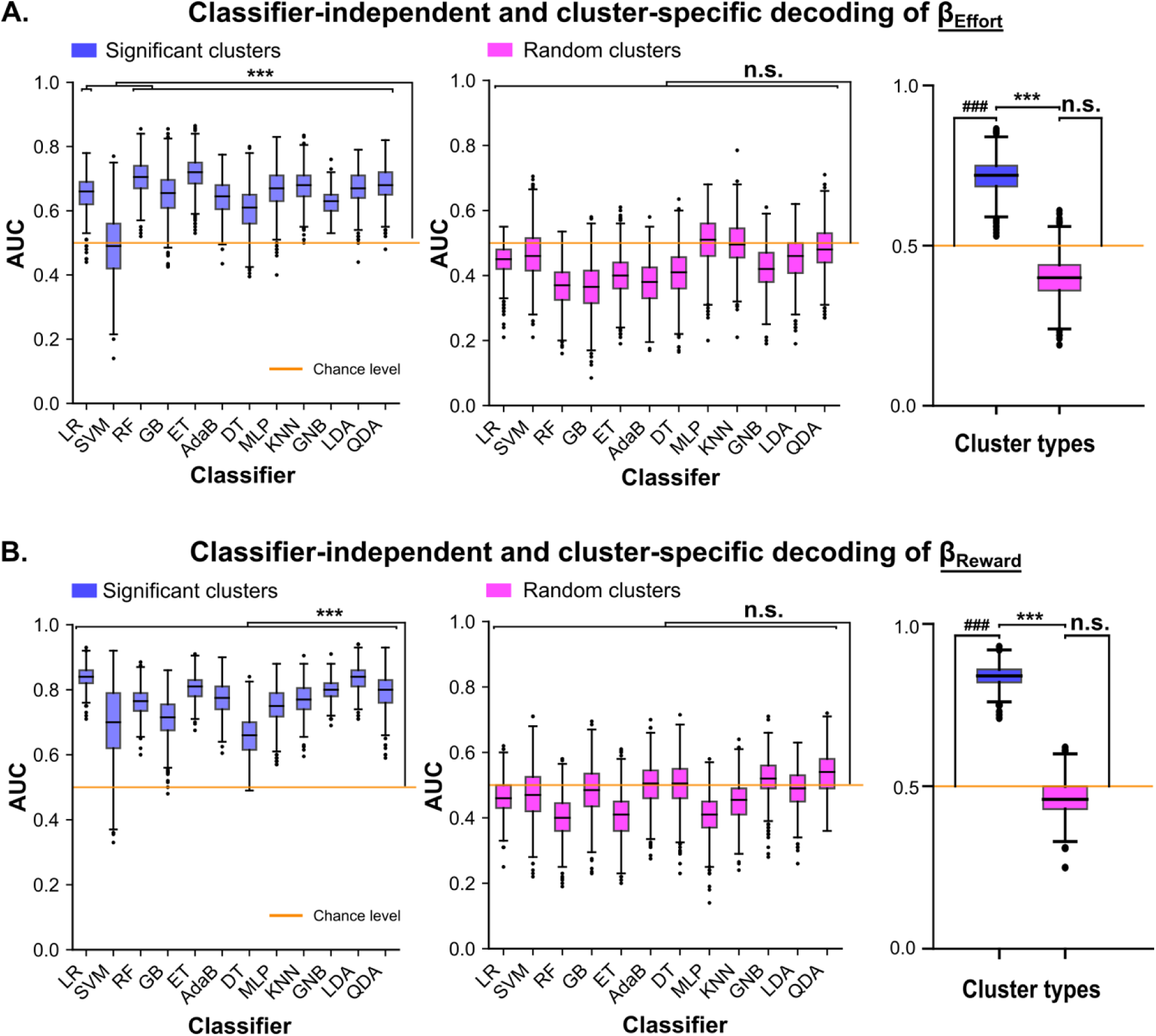
Decoding effort and reward sensitivity is classifier-independent and cluster-specific. **A. Classifier-independent and cluster-specific decoding of β_Effort_.** *Left panel (blue)*: Area under the curve (AUC) values for 12 machine learning classifiers trained on microstructural measures from the 5 clusters significantly associated with β_Effort_. Decoding performance was significantly above chance for nearly all classifiers (all FDR-corrected p < .001), except for SVM (FDR-corrected p = .99), demonstrating classifier-independent decoding. Classifier names: LR: logistic regression; SVM: support vector machine; RF: random forest; GB: gradient boosting; ET: ExtraTrees; AdaB: AdaBoost; DT: decision tree; MLP: multilayer perceptron; KNN: k-nearest neighbors; GNB: Gaussian naive Bayes; LDA: linear discriminant analysis; QDA: quadratic discriminant analysis. *Middle panel (pink)*: AUC values for the same classifiers trained on microstructural measures from randomly sampled white matter clusters, matched in size and number to the original clusters, showing performance near chance. *Right panel*: Statistical comparison of AUC values from the best-performing classifier (ExtraTrees) trained on the 5 significant clusters (blue) versus 5 random clusters (pink). Performance with significant clusters was significantly higher than with random clusters (***FDR-corrected p < .001) and above chance (^###^FDR-corrected p < .001), whereas performance with random clusters did not differ from chance (n.s.). Box plots indicate median and interquartile range; whiskers denote data within 1.5 × IQR. **B. Classifier-independent and cluster-specific decoding of β_Reward_.** The panels follow the same organization as A. but for β_Reward_.

To delineate which white matter clusters most strongly drove the decoding of β_Effort_ and β_Reward_, we conducted feature importance and recursive feature elimination analyses (see *Methods* section). In this context, the term “feature” refers specifically to the individual white matter clusters included as input variables in the classifier, a standard term in machine learning used to denote each parameter or predictor considered during the analysis.

Feature importance rankings (derived from the best-performing classifiers), identified the cluster located within the SMA portion of corticospinal tract as the most predictive feature for both β_Effort_ and β_Reward_ (Figure 10 A and B). This finding aligns with the overlap of this cluster across the β_Effort_ and β_Reward_ cluster-based analyses and its lack of correlation with other computational parameters (*i.e.*, both β_Time_ and β₀), underscoring its specificity to effort and reward sensitivities. Recursive feature elimination indicated that high decoding accuracy could be achieved without the full cluster set. For β_Effort_, classification performance peaked when restricted to the four most informative clusters (AUC = .806 ± .147; Figure 10), three of which were connected to SMA and one with the anterior cingulum. For β_Reward_, classification performance also reached maximal performance with the four most predictive clusters (AUC = .840 ± .136). Of these, two clusters belonged to tracts directly connected to frontal valuation regions (SMA corticospinal tract and OFC-connected forceps minor), while the remaining two involved fronto-parietal and sensorimotor pathways (right superior longitudinal fasciculus and right middle cerebellar peduncle/left cortico-ponto- cerebellar tract). Hence, these analyses indicate that while SMA-connected pathways are the most consistent predictors of both β_Effort_ and β_Reward_, optimal decoding of reward sensitivity additionally relies on contributions from fronto-parietal and motor pathways, reinforcing the idea that inter-individual differences in reward sensitivity emerge from distributed and functionally diverse white matter circuits.

**Figure 10:**
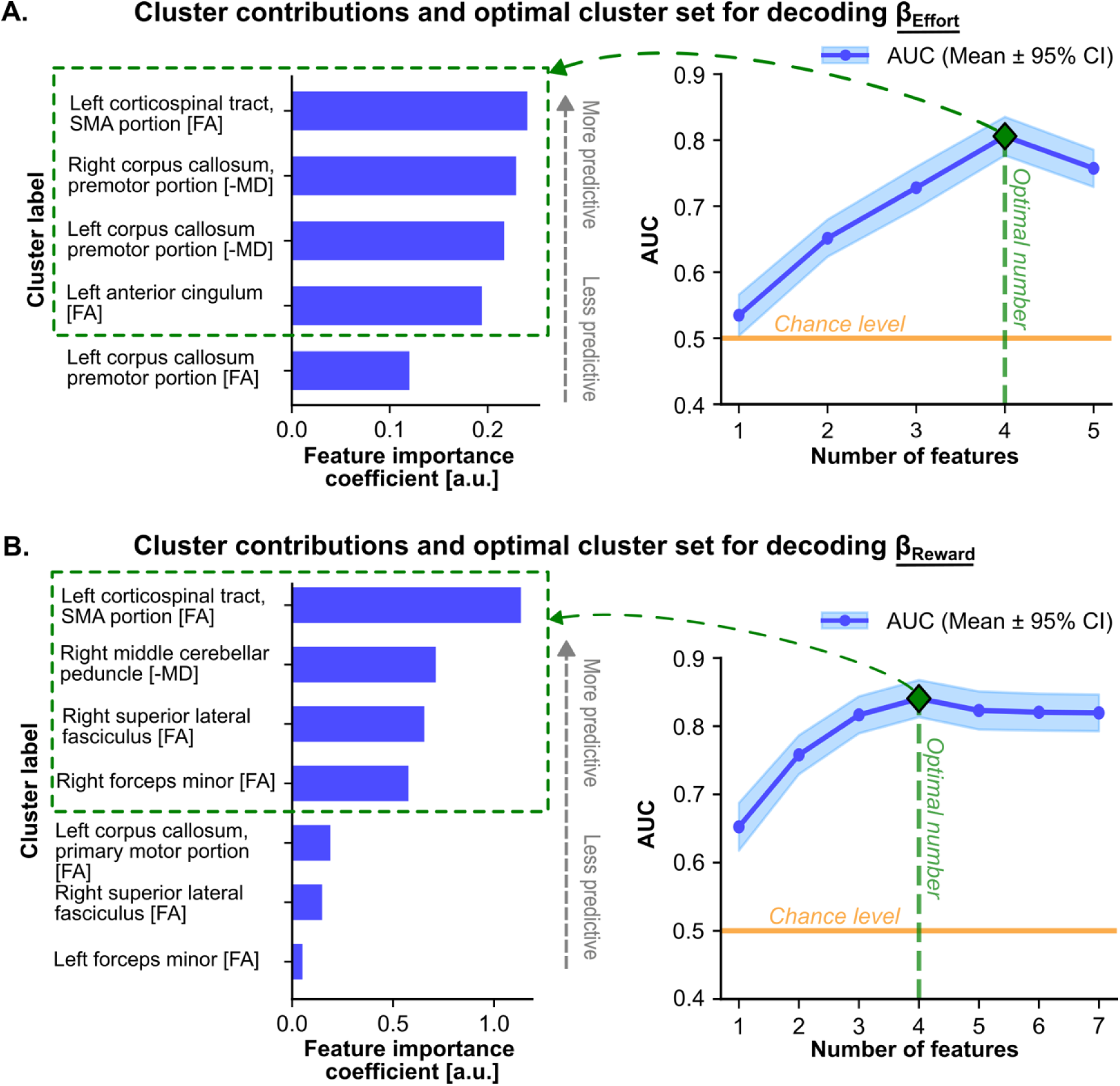
SMA-connected clusters dominate in terms of predictive power, with distributed circuits contributing to reward sensitivity decoding. **A. Cluster contributions and optimal cluster set for decoding β_Effort_.** *Left panel*: Feature importance analysis, where “features” denote the individual white matter clusters used as input variables in the model (a standard machine learning term for predictors). The most predictive cluster was the left corticospinal tract, SMA portion [FA], followed by the right and left premotor portions of the corpus callosum [-MD and FA], and the left anterior cingulum [FA]. *Right panel*: Recursive feature elimination analysis. Classification performance (AUC) peaked when restricted to the four most informative clusters (green diamond), which are highlighted by the green dashed box in the left panel. **A. Cluster contributions and optimal cluster set for decoding β_Reward_.** The panels follow the same organization as A. but for β_Reward_. Here, the left corticospinal tract, SMA portion [FA], was again the most predictive feature, followed by clusters in the cortico-ponto-cerebellar tract [-MD], superior lateral fasciculus [FA], and forceps minor [FA]. Maximal classification performance (AUC) was also reached with four features, indicating that reward sensitivity is best predicted by a combination of SMA-connected, fronto- parietal, and cerebellar pathways.

Altogether, these results demonstrate that white matter microstructure within the clusters identified through voxel-wise analyses carries sufficient information to reliably decode inter-individual differences in both effort and reward sensitivity. Predictions were consistently above chance across complementary metrics (accuracy and AUC) and robust across all tested classifiers. Importantly, classification performance did not exceed chance when using randomly sampled white matter clusters of matched size and number, confirming that predictive information is specific to the loci uncovered by the voxel-wise analyses rather than reflecting generic microstructural variability. Feature importance and recursive elimination further revealed that decoding performance is driven by a subset of clusters, most prominently those connected to SMA, while additional contributions from fronto-parietal and motor pathways are required for optimal prediction of reward sensitivity.

## DISCUSSION

White matter pathways are increasingly recognized as active modulators of neural function, capable of amplifying or attenuating electrophysiological signals (Innocenti et al., 2022; Thiebaut de Schotten and Forkel, 2022), thereby shaping brain activity and behavior (López-Barroso et al., 2013). Variability in white matter microstructure is therefore thought to drive individual differences in cognition and behavior (Forkel et al., 2022), and mapping these structure-behavior relationships can help identify biomarkers for neurological and psychiatric conditions (Thiebaut de Schotten and Forkel, 2022) and predict the behavioral impact of structural lesions (Koch et al., 2021). Recent work highlights the clinical potential of such predictive modelling, from forecasting disease progression (*e.g.*, Parkinson’s, catatonia; Huang et al., 2024, Peretzke et al., 2025) to anticipating behavioral outcomes after neurosurgery (Aylmore et al., 2025; Essayed et al., 2017; Ordonez-Rubiano et al., 2023), suggesting that extending these approaches to motivational constructs could improve risk assessment for apathy and impulsivity and guide personalized interventions. Here, using a whole-brain, data-driven approach combining DWI with computational modelling of decision-making, we show that white matter integrity covaries with individual differences in effort and reward sensitivity, two key determinants of goal-directed behavior. Twelve clusters emerged: 5 linked to effort sensitivity, all within tracts connected to frontal valuation hubs (SMA, dACC, OFC), and 7 linked to reward sensitivity, spanning frontal valuation, fronto-parietal, and sensorimotor pathways. The most robust effects localized to two SMA-connected clusters, one common to effort and reward sensitivity and another converging across FA and -MD metrics and related to effort sensitivity. FA and -MD metrics from the 5 effort-related and 7 reward-related clusters reliably predicted effort and reward sensitivity in out-of-sample machine learning analyses, respectively, whereas randomly sampled clusters did not. SMA-connected tracts dominated decoding in these analyses, but fronto-parietal and sensorimotor pathways also strongly contributed to the decoding of reward sensitivity.

We found both negative and positive associations between white matter integrity and effort/reward sensitivity, supporting the view that white matter modulates neural signals (Innocenti et al., 2022) and that changes in its integrity, whether reductions or increases, alter signal modulation between gray matter regions and influence behavior. Similar bidirectional patterns have been reported across networks in disorders affecting effort and reward sensitivity, including apathy (Baggio et al., 2015; Tay et al., 2019), depression (Leaver et al., 2016; Lynch et al., 2024; Oestreich et al., 2022), and addictions (Tolomeo and Yu, 2022). These observations suggest that both hypo- and hyperconnectivity can heighten sensitivity to effort and reward, depending on the circuit and the stage of dysfunction.

As mentioned above, one of the strongest effects of our data-driven analyses localized to SMA-connected pathways, with a shared cluster in the SMA segment of the corticospinal tract covarying with both effort and reward sensitivity, suggesting a generic role in action valuation. This dual association was construct-specific, as FA values in this cluster did not correlate with β_Time_ and β₀, which index sensitivity to time-on-task and overall baseline acceptance, respectively. Machine learning feature importance analyses independently ranked these SMA clusters as dominant predictors, reinforcing their central role in predicting both effort and reward sensitivity. Although SMA activity has been consistently linked to value computation, particularly in relation to effort processing (Zénon et al., 2015; Bonnelle et al., 2016; Husain and Roiser, 2018; Heron et al., 2019), the prevailing view attributes these changes primarily to interactions with other fronto-striatal valuation hubs, such as the dACC and NAcc (see Husain and Roiser, 2018, for review). Our findings suggest a potentially complementary mechanism: changes in SMA activity may not only reflect interactions with other fronto-striatal valuation regions, but also indicate SMA’s direct influence on behavioral engagement via its corticospinal projections, a pathway largely overlooked in current models. Future studies assessing corticospinal excitability via transcranial magnetic stimulation of the SMA (Entakli et al., 2014; Spieser et al., 2013; Neige et al., 2023) during effort-reward decisions could further characterize the contribution of SMA corticospinal projections to effort and reward sensitivity.

Probabilistic tract labelling further showed that, while the shared cluster indeed mainly overlapped with the SMA corticospinal tract, it also extended beyond this tract, likely engaging additional, more classical valuation-related pathways, such as the SMA-NAcc tract, where reduced macroscopic connectivity has likewise been linked to heightened effort sensitivity (Derosiere et al., 2024). Altogether, reduced integrity in these different SMA-originating circuits may disrupt SMA’s dual roles: that is, invigorating action initiation when movement execution is costly (Fried et al., 1991; Potgieser et al., 2014; Zimnik et al., 2019) and exerting inhibitory control to suppress action initiation when reward incentives are high (Chen et al., 2010), thereby amplifying sensitivity to both effort and reward.

A second major effect involved reduced integrity in the mid-anterior corpus callosum, a key pathway supporting interhemispheric communication, in part between bilateral SMAs, which was robustly associated with heightened effort sensitivity. This effect was strong and spatially consistent, spanning both FA and -MD metrics and forming symmetrical clusters across hemispheres. Extending the view that SMA contributions are not limited to interactions with other frontal valuation nodes (such as the OFC, dACC or NAcc), these findings point to interhemispheric communication as another critical substrate for individual differences in effort computation. Notably, our forceps minor results suggest a parallel role for commissural fibers linking bilateral OFCs in reward sensitivity (see below), underscoring the broader importance of interhemispheric integration in value computation.

In addition to these SMA-connected pathways, additional clusters were identified in classical frontal valuation circuits. As described above, reward sensitivity was positively associated with integrity in the forceps minor, which interconnects bilateral OFCs, regions central to reward processing. Increased connectivity in this tract has also been linked to impulsivity (Jeong et al., 2016), suggesting that stronger OFC communication may enhance reward signal integration and drive greater behavioral engagement when incentives are high. Similarly, higher integrity in the anterior cingulum bundle, connecting SMA and dACC, both key to effort processing, was associated with heightened effort sensitivity. While this might seem at odds with prior findings of reduced cingulum integrity in apathy (Bonnelle et al., 2016), it is consistent with clinical evidence from anterior cingulotomy in obsessive-compulsive disorder, where lesions to this tract reduce action initiation (Bubb et al., 2018). This may suggest that increased integrity in the anterior cingulum might enhance interactions between effort processing regions such as the SMA and dACC (Innocenti et al., 2022; Thiebaut de Schotten and Forkel, 2022), amplifying the influence of high effort costs on behavioral disengagement.

Notably, beyond the canonical frontal valuation regions, our data-driven analysis uncovered less expected white matter pathways linked to reward sensitivity, including the right superior longitudinal fasciculus, cerebellar connections, and the motor segment of the corpus callosum. These associations, unlikely to emerge from hypothesis-driven tractography restricted to predefined valuation circuits, highlight the capacity of whole-brain approaches to reveal non-canonical circuits. While the functional implications remain uncertain, these findings raise the possibility that reward-related signals are transmitted and modulated not only through core valuation hubs but also via broader fronto-parietal and motor networks, potentially influencing how individuals adjust behavioral engagement as rewards increase. Such a role would align with emerging evidence implicating fronto-parietal regions (Etzel et al., 2016), the cerebellum (Kostadinov and Häusser, 2022) and M1 (Derosiere et al., 2017a, 2017b, 2025; Prévost et al., 2010; Pessiglione et al., 2007) in reward-related computations during decision-making.

A key contribution of the present study is the demonstration that individual differences in effort and reward sensitivity can be accurately predicted from white matter microstructure using machine learning. By moving beyond correlational analyses and using a decoding framework, we show that DWI metrics within specific clusters carry sufficient information to decode effort and reward sensitivity across multiple classifiers, highlighting the robustness of these structure-behavior associations. This decoding framework builds on recent advances applying machine learning on DWI metrics in clinical settings (e.g., to predict Parkinson’s progression, Huang et al., 2024 or catatonia, Peretzke et al., 2025) and extends them to dimensional constructs relevant to motivation. Crucially, our findings suggest that microstructure metrics could be leveraged to anticipate the impact of white matter alterations, whether pathological (*e.g.*, following stroke, multiple sclerosis, tumor infiltration, *etc*) or iatrogenic (*e.g.*, following surgical tumor resection, callosotomy, *etc*), on goal-directed behavior. Such predictive modelling may support personalized risk assessment and pre-operative planning in the context of surgery (Aylmore et al., 2025; Essayed et al., 2017; Ordonez-Rubiano et al., 2023), as well as inform targeted rehabilitation strategies (e.g., reward-based interventions, Vassiliadis et al., 2021, 2022), and guide the development of neurotechnological interventions targeting key hubs of the valuation network (Vassiliadis et al., 2024) particularly in cases of motivational impairments such as apathy or impulsivity, which remain challenging to anticipate.

Together, these findings provide a robust anatomical mapping of individual differences in effort and reward sensitivity. By integrating data-driven tract identification with predictive modelling, we isolate specific white matter pathways, particularly SMA-connected tracts, that reliably account for variability in motivational processes. This framework offers a concrete basis for anticipating how structural disruptions may affect goal-directed behavior, with potential relevance for clinical applications in neurology and psychiatry.

## Conflict of interest statement

The authors declare no conflicts of interest

## Acknowledgement

Nam Trinh was supported by the financial grant from Science Foundation Ireland (SFI) Centre for Research Training in Machine Learning at Dublin City University under Grant number 18/CRT/6183. Gerard Derosiere was supported by a FNRS Research associate grant and by an ANR-JCJC grant (ANR-24-CE37-6839-01). Tomás Ward is supported by Research Ireland under grant agreement 12/RC/2289_P2. Pierre Vassiliadis was supported by a Swiss National Science Foundation (SNSF) fellowship (P500PB_230720). Julie Duque was supported by three FNRS research grants (T008219F, J005921F and T007023F). We would like to thank Matthieu Boisgontier, Ahmad Noureddine and Valentin Touzé for their participation in data acquisition.

## AUTHOR CONTRIBUTIONS

NT: Machine learning analysis, interpretation of the results, data visualisation, article writing. LD, QD: MRI data processing, article writing. PV: computational modelling, interpretation of the results, article writing. JD: study conception, interpretation of the results, article writing. TW: Machine learning analysis, interpretation of the results. GD: study conception, data acquisition, interpretation of the results, article writing from the first draft to the final version.

## Notes

### Competing Interest Statement

The authors have declared no competing interest.

