## Supplementary materials for "White matter microstructure predicts effort and reward sensitivity"

**A.** **Candidate models**

1. $Value = \beta_{Effort} \times EFFORT_{LEVEL} + \beta_{Reward} \times REWARD_{LEVEL} + \beta_0$

2. $Value = \beta_{Effort} \times EFFORT_{LEVEL} \times (1 + \beta_{Time} \times t) + \beta_{Reward} \times REWARD_{LEVEL} + \beta_0$

3. $Value = \beta_{Effort} \times EFFORT_{LEVEL}{}^2 + \beta_{Reward} \times REWARD_{LEVEL} + \beta_0$

4. $Value = \beta_{Effort} \times EFFORT_{LEVEL}{}^2 \times (1 + \beta_{Time} \times t) + \beta_{Reward} \times REWARD_{LEVEL} + \beta_0$

5. $Value = e^{\beta_{Effort} \times EFFORT_{LEVEL}} + \beta_{Reward} \times REWARD_{LEVEL} + \beta_0$

6. $Value = e^{\beta_{Effort} \times EFFORT_{LEVEL} \times (1+\beta_{Time} \times t)} + \beta_{Reward} \times REWARD_{LEVEL} + \beta_0$

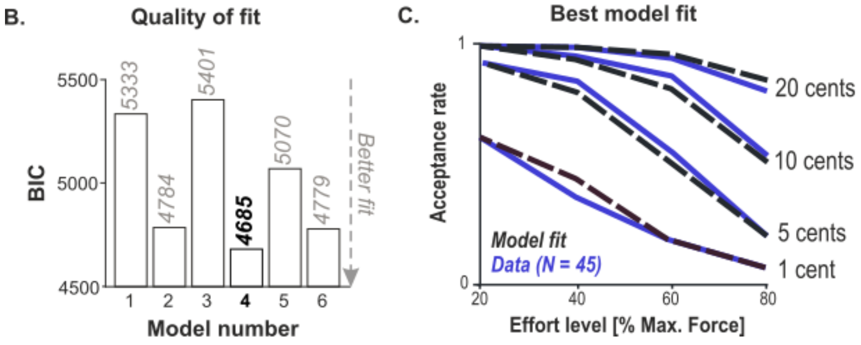


**Supplementary Figure 1: Computational modelling of effort-based decision-making. A. Candidate models.** Candidate models of subjective value computation were tested to quantify individual differences in effort and reward sensitivity. Models differed in cost function (linear, quadratic, or exponential) and in whether they included a time-dependent modulation term ($\beta_{Time}$), following procedures established in prior work (e.g., (Le Heron et al., 2018a)). Value was modelled as a function of effort level ($EFFORT_{LEVEL}$), reward level ($REWARD_{LEVEL}$), and trial number (t), with free parameters capturing individual sensitivities ($\beta_{Effort}$, $\beta_{Reward}$) and baseline bias ($\beta_0$). **B. Model fit comparison based on Bayesian information criterion (BIC)**. Model 4, which combined a quadratic cost function with a time-modulated cost term, showed the best fit (lowest BIC), consistent with prior findings (Le Heron et al., 2018b; Prévost et al., 2010; Pessiglione et al., 2018). **C. Best-fitting model vs. observed behavior.** Model-derived acceptance rate (black dashed lines) closely captured observed behavioral data (blue lines; N = 45), across varying effort levels and reward magnitudes.

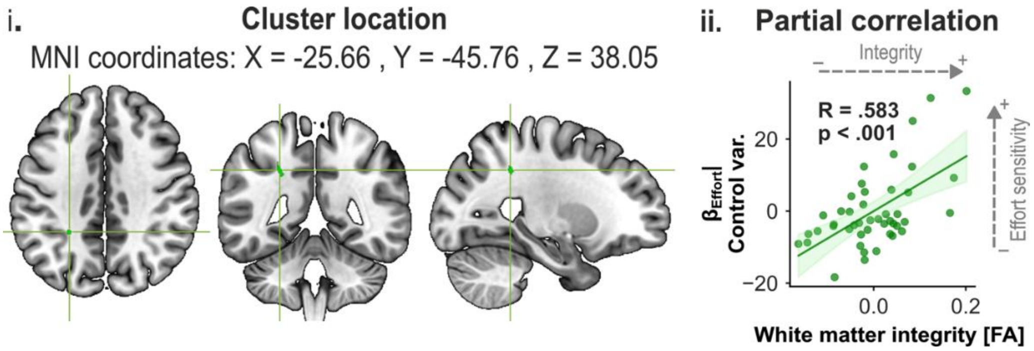


**Supplementary Figure 2. Superficial cluster showing a positive association between fractional anisotropy (FA) and $\beta_{Effort}$.** i. A significant cluster was identified outside canonical white matter tracts, located in the left parietal cortex (MNI coordinates: X = −25.66, Y = −45.76, Z = 38.05; volume: 62 mm³). ii. Partial correlation analysis confirmed this association, revealing a robust positive correlation between FA values extracted from the cluster and $\beta_{Effort}$ ($R = .583$, $p < .001$). These findings suggest that, in addition to the major tracts identified using a probabilistic white matter atlas, more superficial white matter regions may also contribute to individual variability in effort sensitivity. This further highlights the added value of whole-brain, data-driven approaches for uncovering relevant structural substrates beyond predefined tracts-of-interest.

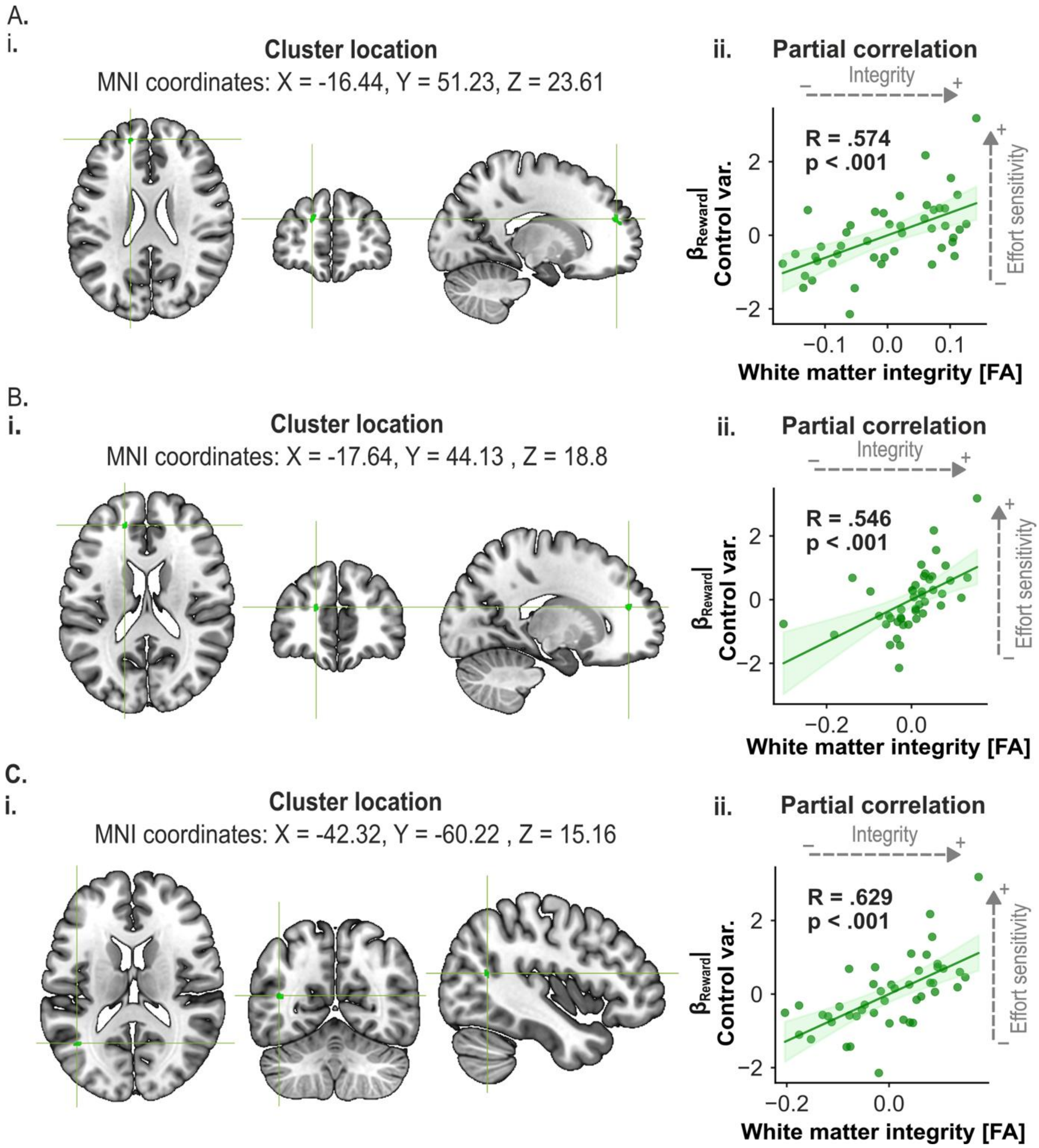


**Supplementary Figure 3: Three clusters with positive $\beta_{Reward}$ associations in superficial white matter of the left frontal and parietal cortex outside major tracts.** Three additional clusters exhibiting significant positive associations between white matter integrity and reward sensitivity ($\beta_{Reward}$) were identified in superficial regions of the left frontal and parietal cortex, outside canonical major white matter tracts. **A. i.** Cluster (green) located at MNI coordinates: X = −16.44, Y = 51.23, Z = 23.61; volume: 88 mm³. **ii.** Partial correlation between fractional anisotropy (FA) in this cluster and $\beta_{Reward}$: $R$ = .574, $p$ < .001. **B. i.** Cluster (green) located at MNI coordinates: X = −17.64, Y = 44.13, Z = 18.88; volume: 64 mm³. ii. Partial correlation with $\beta_{Reward}$: $R$ = .546, $p$ < .001. **C. i**. Cluster (green) located at MNI coordinates: X = −42.32, Y = −60.22, Z = 15.16; volume: 64 mm³). **ii.** Partial correlation with $\beta_{Reward}$: $R$ = .629, $p$ < .001.

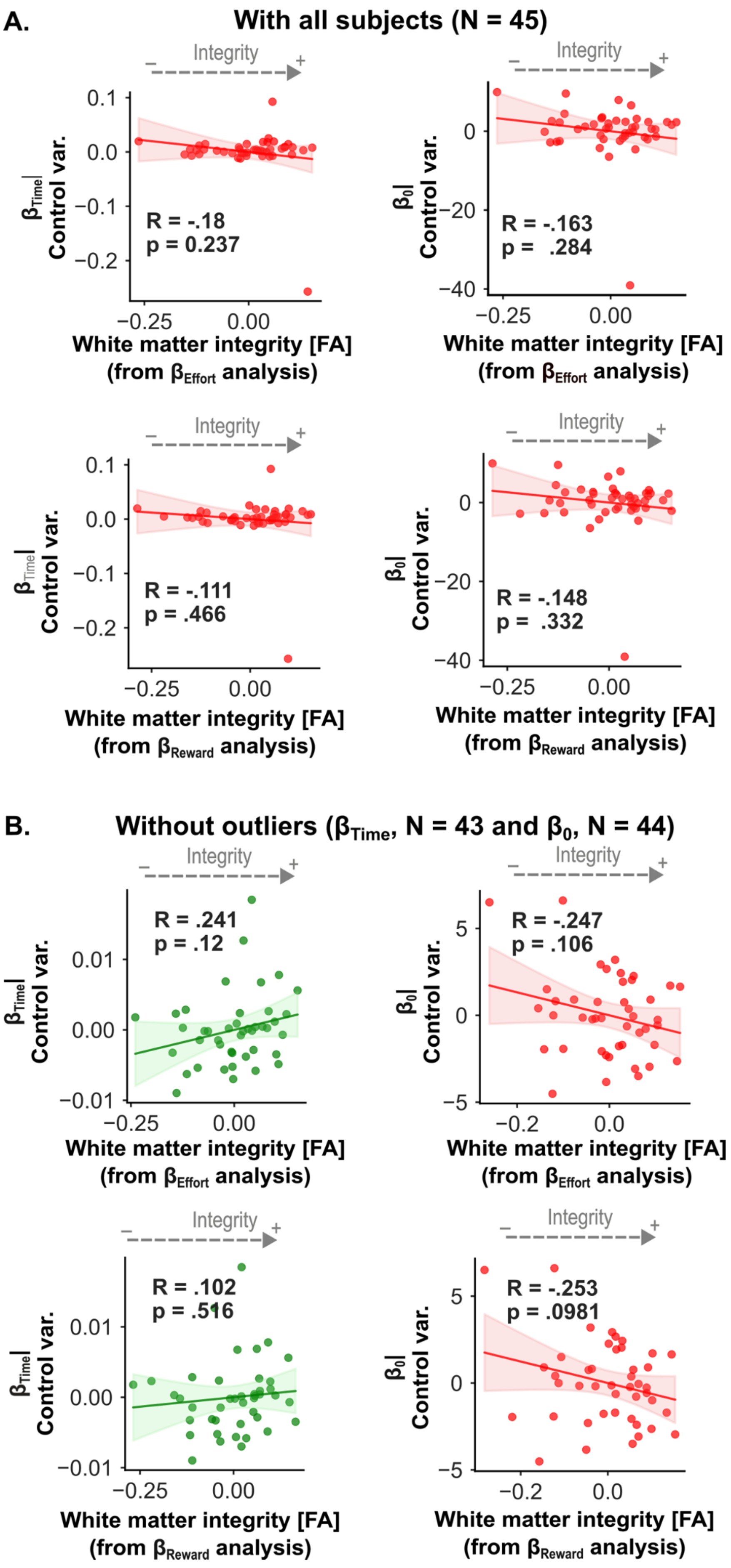
A.
With all subjects (N = 45)
Integrity
R = -.18
p = 0.237
R = -.163
p = .284
R = -.111
p = .466
R = -.148
p = .332
$\beta_{Time}$| Control var.
$\beta_0$| Control var.
White matter integrity [FA] (from $\beta_{Effort}$ analysis)
White matter integrity [FA] (from $\beta_{Reward}$ analysis)
B.
Without outliers ($\beta_{Time}$, N = 43 and $\beta_0$, N = 44)
R = .241
p = .12
R = -.247
p = .106
R = .102
p = .516
R = -.253
p = .0981

**Supplementary Figure 4: The SMA cluster associated with both $\beta_{Effort}$ and $\beta_{Reward}$ is specific and unrelated to the two other model parameters.** To confirm that the dual association with $\beta_{Effort}$ and $\beta_{Reward}$ was not trivially driven by their derivation from the same acceptance rates, we tested whether FA values in these SMA clusters correlated with the two other model-derived parameters, $\beta_{Time}$ and $\beta_0$. **A. Correlations including all participants (n = 45).** No significant correlations were observed for either the effort-related cluster ($\beta_{Time}$: R = -.180, p = .237; $\beta_0$: R = -.163, p = .284) or the reward-related cluster ($\beta_{Time}$: R = -.111, p = .466; $\beta_0$: R = -.148, p = .332), underscoring the specificity of this SMA white matter locus for effort and reward sensitivities. B. Correlations excluding outliers in $\beta_{Time}$ and $\beta_0$. Results remained consistent, with no significant correlations for either cluster (effort-related: $\beta_{Time}$: R = .241, p = .120; $\beta_0$: R = -.247, p = .106; reward-related: $\beta_{Time}$: R = .102, p = .516; $\beta_0$: R = -.253, p = .0981). Across analyses, FA values in these clusters did not relate to $\beta_{Time}$ or $\beta_0$, confirming that this SMA cluster specifically covaries with individual differences in effort- and reward-related parameters.

| Model | AUC | Accuracy | Recall | Precision | F1 |
|---|---|---|---|---|---|
| LR | 0.652 ± 0.053 | 0.613 ± 0.042 | 0.580 ± 0.063 | 0.644 ± 0.075 | 0.586 ± 0.058 |
| SVM | 0.490 ± 0.099 | 0.561 ± 0.055 | 0.484 ± 0.096 | 0.590 ± 0.109 | 0.497 ± 0.085 |
| RF | 0.705 ± 0.052 | 0.630 ± 0.054 | 0.612 ± 0.060 | 0.665 ± 0.080 | 0.613 ± 0.056 |
| GB | 0.651 ± 0.068 | 0.601 ± 0.060 | 0.603 ± 0.080 | 0.627 ± 0.081 | 0.592 ± 0.067 |
| **ET** | **0.719 ± 0.050** | **0.648 ± 0.053** | **0.602 ± 0.064** | **0.698 ± 0.084** | **0.619 ± 0.059** |
| AdaB | 0.644 ± 0.055 | 0.568 ± 0.051 | 0.601 ± 0.079 | 0.588 ± 0.072 | 0.568 ± 0.061 |
| DT | 0.606 ± 0.067 | 0.592 ± 0.064 | 0.588 ± 0.088 | 0.620 ± 0.090 | 0.579 ± 0.073 |
| MLP | 0.670 ± 0.061 | 0.605 ± 0.053 | 0.597 ± 0.075 | 0.633 ± 0.072 | 0.592 ± 0.062 |
| KNN | 0.677 ± 0.049 | 0.604 ± 0.045 | 0.664 ± 0.054 | 0.618 ± 0.055 | 0.624 ± 0.044 |
| GNB | 0.625 ± 0.034 | 0.572 ± 0.035 | 0.386 ± 0.044 | 0.627 ± 0.103 | 0.449 ± 0.048 |
| LDA | 0.671 ± 0.048 | 0.634 ± 0.037 | 0.568 ± 0.049 | 0.679 ± 0.065 | 0.596 ± 0.048 |
| QDA | 0.682 ± 0.055 | 0.612 ± 0.046 | 0.430 ± 0.062 | 0.698 ± 0.115 | 0.500 ± 0.065 |

**Supplementary Table 1: Decoding effort sensitivity is classifier-independent: further evidence from multiple machine learning metrics.** Performance of 12 machine learning classifiers in predicting effort sensitivity from white matter microstructural features, including fractional anisotropy and mean diffusivity. Reported values reflect the mean ± standard deviation across 1,000 iterations of stratified 5-fold cross-validation with randomized data partitioning. Classifier abbreviations: LR, logistic regression; SVM, support vector machine; RF, random forest; GB, gradient boosting; ET, extra trees; AdaB, AdaBoost; DT, decision tree; MLP, multi-layer perceptron; KNN, k-nearest neighbors; GNB, Gaussian naïve Bayes; LDA, linear discriminant analysis; QDA, quadratic discriminant analysis. Performance metrics include area under the receiver operating characteristic curve (AUC), accuracy, recall (sensitivity), precision (positive predictive value), and F1 score (harmonic mean of precision and recall). Among classifiers, the ET model achieved the highest AUC (0.719 ± 0.050), along with a balance between precision (0.698 ± 0.084) and F1 score (0.619 ± 0.059), indicating robust discriminative performance. RF and QDA also performed well (AUCs of 0.705 ± 0.052 and 0.682 ± 0.055, respectively), though QDA exhibited reduced recall (0.430 ± 0.062), suggesting a trade-off favoring specificity. Linear models such as LR and LDA demonstrated moderate AUCs (0.652 ± 0.053 and 0.671 ± 0.048, respectively), with LDA yielding relatively high precision (0.679 ± 0.065). In contrast, SVM showed the lowest AUC (0.490 ± 0.099), indicating performance at or below chance levels. GNB and QDA showed the largest discrepancies between precision and recall, highlighting imbalances in classification. Hence, despite variations in individual metrics, most classifiers achieved above-chance performance, reinforcing the robustness of the decoding. These results collectively indicate that effort sensitivity can be reliably predicted from white matter features across a wide range of machine learning approaches.

| Model | AUC | Accuracy | Recall | Precision | F1 |
|---|---|---|---|---|---|
| **LR** | **0.837 ± 0.034** | **0.718 ± 0.038** | **0.749 ± 0.052** | **0.736 ± 0.047** | **0.722 ± 0.043** |
| SVM | 0.695 ± 0.117 | 0.704 ± 0.041 | 0.787 ± 0.064 | 0.707 ± 0.049 | 0.721 ± 0.046 |
| RF | 0.763 ± 0.042 | 0.678 ± 0.039 | 0.689 ± 0.059 | 0.702 ± 0.053 | 0.673 ± 0.047 |
| GB | 0.711 ± 0.058 | 0.675 ± 0.054 | 0.702 ± 0.083 | 0.691 ± 0.064 | 0.674 ± 0.067 |
| ET | 0.807 ± 0.037 | 0.693 ± 0.040 | 0.743 ± 0.057 | 0.707 ± 0.051 | 0.702 ± 0.045 |
| AdaB | 0.774 ± 0.048 | 0.710 ± 0.051 | 0.773 ± 0.073 | 0.712 ± 0.060 | 0.720 ± 0.058 |
| DT | 0.656 ± 0.059 | 0.632 ± 0.055 | 0.640 ± 0.092 | 0.650 ± 0.071 | 0.620 ± 0.071 |
| MLP | 0.750 ± 0.052 | 0.674 ± 0.052 | 0.685 ± 0.070 | 0.694 ± 0.068 | 0.668 ± 0.056 |
| KNN | 0.771 ± 0.045 | 0.685 ± 0.045 | 0.731 ± 0.058 | 0.699 ± 0.053 | 0.694 ± 0.048 |
| GNB | 0.802 ± 0.030 | 0.697 ± 0.025 | 0.815 ± 0.042 | 0.682 ± 0.026 | 0.729 ± 0.029 |
| LDA | 0.834 ± 0.035 | 0.721 ± 0.036 | 0.771 ± 0.047 | 0.731 ± 0.044 | 0.732 ± 0.039 |
| QDA | 0.792 ± 0.049 | 0.692 ± 0.044 | 0.807 ± 0.065 | 0.683 ± 0.045 | 0.722 ± 0.047 |

**Supplementary Table 2: Decoding reward sensitivity is classifier-independent: further evidence from multiple machine learning metrics.** Performance of 12 machine learning classifiers in predicting reward sensitivity from white matter microstructural features, including fractional anisotropy and mean diffusivity. Reported values reflect the mean ± standard deviation across 1,000 iterations of stratified 5-fold cross-validation with randomized data partitioning. Classifier abbreviations: LR, logistic regression; SVM, support vector machine; RF, random forest; GB, gradient boosting; ET, extra trees; AdaB, AdaBoost; DT, decision tree; MLP, multi-layer perceptron; KNN, k-nearest neighbors; GNB, Gaussian naïve Bayes; LDA, linear discriminant analysis; QDA, quadratic discriminant analysis. Performance metrics include area under the receiver operating characteristic curve (AUC), accuracy, recall (sensitivity), precision (positive predictive value), and F1 score (harmonic mean of precision and recall). Across models, LR and LDA yielded the highest AUC values (0.837 ± 0.034, highlighted in the table and 0.834 ± 0.035, respectively), indicating strong overall discriminative ability. GNB and QDA also performed competitively (AUCs of 0.802 ± 0.030 and 0.792 ± 0.049, respectively), with GNB achieving the highest recall (0.815 ± 0.042), suggesting strong sensitivity to effort-related signal in the input features. In contrast, tree-based methods such as DT and GB exhibited comparatively lower performance across most metrics, highlighting limitations in their generalizability to the underlying data structure. While MLP and KNN classifiers offered moderate performance, ensemble methods like AdaB and ET demonstrated a balance between precision and recall, though their AUCs fell below those of the linear models. These results suggest that linear decision boundaries may be particularly well suited for classifying reward sensitivity from microstructural white matter features, potentially reflecting a low-dimensional organization of the relevant neural representations. Further, despite differences in classifier type and complexity, most models achieved consistent above-chance accuracy, demonstrating that reward sensitivity is robustly decodable. These results again indicate that reward sensitivity can be reliably predicted from white matter features across a wide range of machine learning approaches.

**Supplementary material references**

Le Heron C, Apps. MAJ, Husain M (2018a) The anatomy of apathy: A neurocognitive framework for amotivated behaviour. Neuropsychologia 118:54–67.

Le Heron C, Manohar S, Plant O, Muhammed K, Griffanti L, Nemeth A, Douaud G, Markus HS, Husain M (2018b) Dysfunctional effort-based decision-making underlies apathy in genetic cerebral small vessel disease. Brain 141:3193–3210.

Pessiglione M, Vinckier F, Bouret S, Daunizeau J, Le Bouc R (2018) Why not try harder? Computational approach to motivation deficits in neuro-psychiatric diseases. Brain 141:629–650.

Prévost C, Pessiglione M, Météreau E, Cléry-Melin M-L, Dreher J-C (2010) Separate Valuation Subsystems for Delay and Effort Decision Costs. J Neurosci 30:14080–14090.
